# Investigating cognitive enrichment for dairy calves through behavioral measures of participation and engagement: a pilot study

**DOI:** 10.64898/2026.04.01.715895

**Authors:** Georgiana Amarioarei, Marjorie Cellier, Nadège Aigueperse, Tania Wolfe, Elise Shepley, Abdoulaye Baniré Diallo, Elsa Vasseur

## Abstract

Introducing cognitive enrichment from an early age has the potential to enhance an animal’s capacity to learn both simple and complex tasks, promote neural plasticity, and support cognitive development. This is applicable for young cattle who are at a critical stage in their development and could benefit from the influence cognitive enrichment has on their behavioral expression. This study aims to explore the effects cognitive enrichment has on weaned dairy calves through analyzing behavioral measures of voluntary participation and short-term behavioral reactions to enrichment exposure. Our study involved a total of five pairs of weaned calves (n=8 treatment; n=2 control). The treatment groups were presented with three variations of a puzzle box, each equipped with unique challenges that offer different solutions (push, slide, pull). These boxes were provided to the calves twice daily over the span of nine days in an isolated corridor located behind their pen. We hypothesized that motivated calves would consistently engage with cognitive enrichment voluntarily over time and express directed natural behaviors, reflecting sustained participation across repeated trials. Results demonstrated that calves consistently visited the cognitive enrichment area across trials, with an average latency of 75.7 ± 47.0s from the pen to the enrichment. Secondly, the calves spent a significant proportion of trial time within the enrichment area at 65% (870.1 ± 21s). Lastly, all calves expressed a broad range of behaviors in line with their natural exploration within the enrichment area, while the puzzle box treatment groups expressed higher durations of behavioral expressions when compared to the control (F=11.7*, p*<0.0001). Combined, these results indicate the calves’ motivations to voluntarily participate in a cognitive challenge. While these are promising findings for cognitive enrichment and its applicability to dairy calves, further work is needed to understand broader parameters. Specifically, how can social dynamics influence enrichment interaction in groups, how can this type of enrichment be implemented on farms, and what are the long-term effects to providing cognitive enrichment in the early stages of development.

## 1. INTRODUCTION

In contrast to their wild counterparts, animals in captivity tend to live in highly predictable and structured environments where they are infrequently or inappropriately challenged (Morgan & Tromborg, 2007; Wemelsfelder & Birke, 1997). Over the last decades, attention to animal welfare has increased globally as a result of continued intensification of livestock production systems, technological innovations, evolving dietary habits, and changes in consumer perception (Alonso et al., 2020; Broom, 2022). The source of some of these welfare concerns are rooted in the minimized environmental stimulation animals are receiving from their standardized and unnatural environments. While highly efficient for monitoring and management, intensive housing may lead to chronic boredom which in the long term may create health, performance and behavioral problems (Wemelsfelder, 1993, Mason et al., 2007; Veissier et al., 2024). In addition to boredom, animals housed in barren or restrictive housing are unable to perform many of their species-specific behaviors which have been noted as a major source of stress leading to impairment of health and overall welfare (Morgan & Tromborg, 2007). Consequently, initiatives to improve animal care and their respective environments have been made with the goal to satisfy some of the ethical implications associated with animal production systems (Alonso et al., 2020). One of these improvements includes providing augmentations or alterations to the environment in the form of enrichment. Environmental enrichment is defined by Shepherdson (1998) as an animal husbandry principle that seeks to enhance the quality of captive animal care by identifying and providing the environmental stimulus necessary for optimal psychological and physiological well-being. Supplying animals living in intensive housing with some forms of enrichment can provide the opportunity to explore a more sophisticated environment, gain more active control, and reduce boredom and its negative consequences on well-being, health and behavior. (Manteuffel et al., 2009).

On average, animals are highly motivated to explore and acquire resources under a variety of conditions, even when resources and necessities are concurrently available with little or no effort on the animal’s part (Wemelsfelder & Birke, 1997). Motivation in animals can be defined as the internal drive of an animal to perform behavior as a result of their perceived physiological or psychological state and studies point towards the importance of improving systems to allow animals to act on these motivations (Jensen & Toates, 1993; Manteuffel et al., 2009; Muszik, 2025). Research on animal motivation suggests that animals benefit from and may even prefer a more complex environment that allows for exploration and positive stimuli (Jensen & Toates, 1993; Morgan & Tromborg, 2007; Clark, 2017). While a complex environment is not always feasible within a commercial setting, environmental enrichment is an acceptable alternative that can provide interactive objects that reap similar benefits. Specifically, animals reared in enriched housing are often less reactive, quicker to acquire tasks and better able to adapt to changes than animals reared in perceptually poor environments (Zhang et al., 2022). However, in order to properly benefit from an effective enrichment, a criterion must first be met. A successful environmental enrichment should provide the animal with more control over their environment, promote natural behavioral expression, support species-appropriate repertoires and allow the animal to adequately deal with challenges (Mench, 1998; Veissier et al., 2024). Lastly, the enrichment should allow for sustained engagement or be routinely modified/replaced to maintain the animal’s interest.

Cognitive enrichment is a subset of environmental enrichment that refers to the structured delivery of cognitively engaging opportunities that promote goal-directed behaviors, operant learning, and perceptual discrimination in animals, thereby engaging evolved cognitive capacities such as problem-solving, memory, and environmental assessment (Manteuffel et al., 2009; Zebunke et al., 2013; Kleiber et al., 2023). By enabling operant learning and engagement with the use of meaningful rewards such as food, water, social contact, cognitive enrichment allows animals to exert agency, cope with their surroundings, and express natural behaviors. Cognitive enrichment is unique from other environmental enrichments because it is intentionally designed to target specific cognitive processes (like memory and learning) and aims to measure the otherwise imperceptible process of cognitive stimulation (Shettleworth, 2010; Clark, 2017). It is crucial that cognitive enrichment matches the level of challenge an animal requires for their cognitive skill level such that the animal can be occupationally and psychologically satisfied (Wemelsfelder & Birke, 1997; Kleiber et al., 2023). Specifically, simple puzzles can become rapidly disinteresting without a mechanism to vary the challenge, whereas puzzles of high difficulty can result in frustration (Meehan & Mench, 2007). Thus, if the cognitive enrichment is not a good match for the animal, its purpose is nullified, and the animal will not experience the intended positive effects. Past research on the link between cognitive challenge and well-being has highlighted that a lack of cognitive challenge in captive environments is, at best, a missed opportunity to increase welfare, and at worst, a source of negative welfare (Meehan & Mench, 2007).

While its application remains relatively uncommon, cognitive enrichment has been broadly employed in species considered to possess advanced intelligence such as great apes (Clark, 2011; Morimura, 2006), elephants (Foerder et al., 2011) and cetaceans (Harley et al., 2010). However, emerging research demonstrates that taxa long assumed to be cognitively simple exhibit complex cognitive abilities and may also derive benefits from opportunities to exercise them (Hagen & Broom, 2004; Zentall, 2021). Farm animal species in particular have well-developed sensory and cognitive abilities (Croney et al., 2003; Boissy et al., 2007) and could benefit from the positive welfare outcomes associated with cognitive enrichment. Furthermore, introducing cognitive enrichment from a younger age in the animal’s life has been studied to support wellbeing, promote neural plasticity, and support cognitive development (Zentall, 2021; Salvanes et al., 2013). Younger animals are more susceptible to developing new behaviors resulting from cognitive activity than older ones (Milgram 2003). In turn, cognitive enrichment can support coping behaviors for developing animals that are useful through adulthood. This is especially relevant for populations that deal with stressful situations such as displacement and environmental change. One such animal that could benefit from the implementation of cognitive enrichment is a dairy calf. Calves are at a critical stage in their development and often experience situations where they need to adapt quickly to environmental change as they age. To experience their environments, calves express different types of non-nutritive oral behaviors such as manipulating substrates of their home pen in the form of licking, nibbling, biting or suckling (Le Neindre, 1993). Sometimes these behaviors are perceived as negative because they can be damaging to the housing materials, but these behaviors also serve as learning tools that are key to their neural processing. Consequently, cognitive enrichment has the potential to redirect natural exploratory behaviors while supporting the learning abilities and coping skills of calves.

While research has been conducted on cognitive abilities of calves, to the best of our knowledge, no studies exploring cognitive enrichment for calves have been published to date. While similar in name, cognitive tests and cognitive enrichment differ significantly from each other. Firstly, the goals are different in that a cognition test focuses on evaluating cognitive processes, abilities and performances and requires repeated testing of known individuals under standardized conditions in a way that allows for noise caused by differences to be identified, quantified and/or removed (Lauber et al., 2006; Gaillard et al., 2014; Clark, 2017). On the other hand, cognitive enrichment tends to have looser parameters that focus less on the outcome of the animal’s performance but more on how the enrichment influences the animal’s time budgets, behaviors and motivations in the short and long term.

The main objective of this animal study is to investigate the effects cognitive enrichment has on weaned dairy calves through analyzing behavioral measures of voluntary participation and short-term behavioral reactions to enrichment exposure. The cognitive enrichment used consisted of three puzzle boxes with varying solutions of intended equivalent difficulty provided on a randomized and unpredictable rotation over the course of nine days. We hypothesize that motivated calves will consistently engage with cognitive enrichment voluntarily over time and express directed natural behaviors, reflecting sustained participation across repeated trials. We predict that at the pen level, calves will choose to spend more time in the enrichment area than remaining inside the pen. Secondly, we predict latencies to visit the enrichment and time spent interacting will either increase or remain consistent over time. Thirdly, solving the box ability will not influence the interaction levels of calves. Lastly, if calves are motivated to interact with the cognitive enrichment we expect a diverse repertoire of exploratory behaviors across repeated trials. This project intends to serve as a pilot study to begin understanding how calves react to and utilize cognitive enrichment when the opportunity is available.

## 2. MATERIALS AND METHODS

### 2.1. Ethical Statement

All research procedures and the use of animals were approved by the Animal Care Committee of McGill University and affiliated hospitals and research institutes (Protocol #MCGL-10059).

### 2.2. Animals and Experimental Design

Ten female Holstein calves were paired together (5 pairs) in double pens (5.5 x 3m) based on birth date and as pairs, were weaned and dehorned at the same time. Each pair of calves was enrolled in the experiment a month after weaning and two months after disbudding (see Figure 1. for enrollment details). Selection criteria was restricted to female Holsteins that were in good health and had no complications from the dehorning process that would have affected the time of enrollment. The study took place intermittently from September to December 2024 and the approximate age of the calves at the start of the experiment was 4 months (± 2 weeks). The calves were housed indoors in a heated calf barn at the Macdonald Campus Dairy Cattle Complex of McGill University. The pens were organized in two rows of four with calves from adjacent pens having visual, auditory and limited physical contact. Non-adjacent pens had visual and auditory but no physical contact with one another. The pens were equipped with a straw bedded laying area, brush enrichment and a water trough. All calves had *ad-libitum* access to water and were fed twice daily (morning and evening).

**Figure 1.**
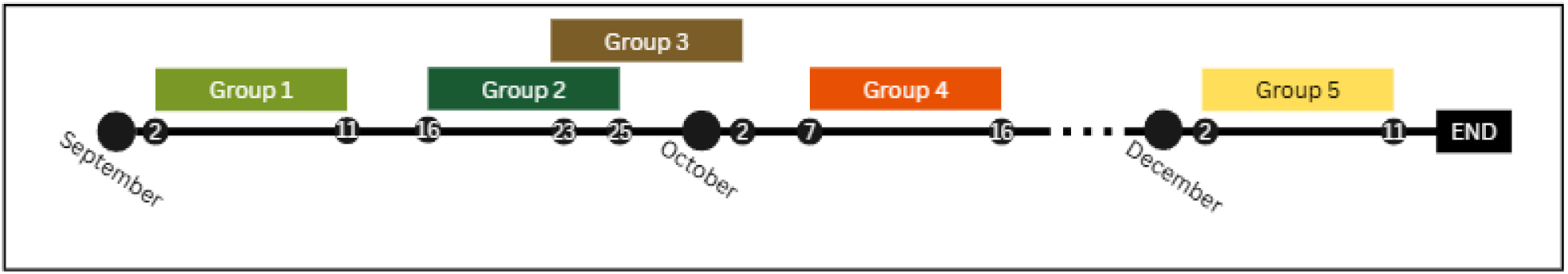
Enrollment timeline of calf groups for the experimental period. Intermittently enrolled based on age and number of weeks post dehorning.

### 2.3. Pre-Experimental Phase

#### 2.3.1. Food Reward Habituation

Two weeks postweaning, calves were habituated to the novel food reward through a three-day exposure process (see Figure 2). The food selected as the reward for the experiment was a mixture of locally acquired fresh green cabbage, bananas and red seedless watermelon. This reward was justified through the consultation of a registered practicing veterinarian (Macdonald Dairy Cattle Complex herd veterinarian), previous research (Detering, 1976; Mukodiningsih et al., 2017), and the results of a supplementary preference test conducted on non-experiment calves.

**Figure 2.**
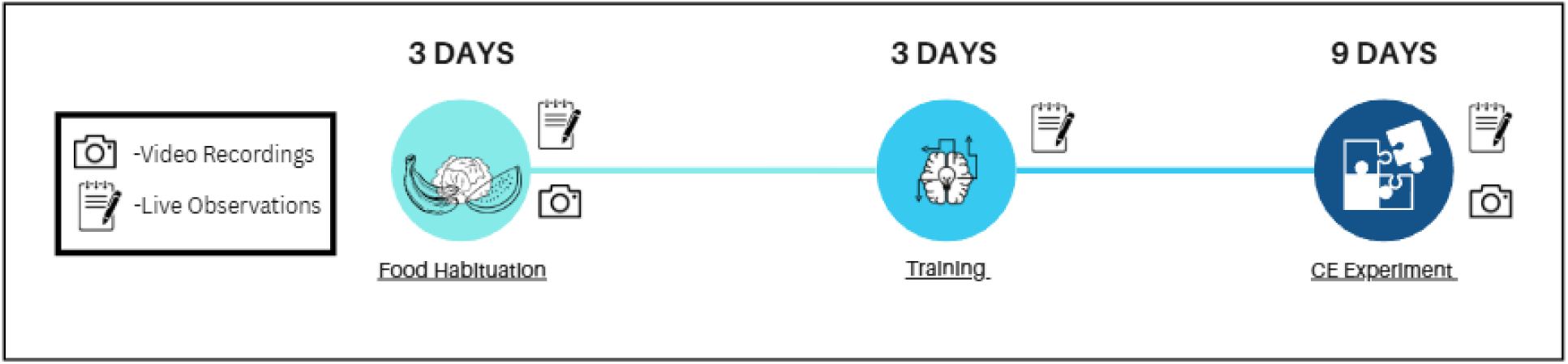
Timeline of calf food habituation, training period and cognitive enrichment (CE) experiment durations. During each phase, data was recorded in the form of live observations. Additionally, for food habituation and the CE experiment, video recordings were taken for later analysis.

The food rewards were kept unmixed and about ½ cup of each (125ml) was divided into identical purple feed buckets (Fortiflex©: 16qt Hook over feeder, San Juan, Puerto Rico), one per food option (3 total). The feed buckets were then placed in the food alley in front of the calf pen in a randomized order and a timer for 15 minutes began once the researchers had exited the barn. A GoPro HERO11 camera (GoPro Inc., San Mateo, CA, USA) mounted on a tripod in front of the pen recorded the behaviors of the calves during this time. After 15 minutes, there was a 30-minute pause in which the buckets were removed, then returned for a second 15-minute sessions following the same procedure as previously stated. This process was repeated for the following two days (three total) such that the calves received overall six food exposures in randomized orders to avoid associating bucket positions with a particular food. Calves passed the food habituation phase if both individuals within the pen consumed the food from the buckets at least once over the three-day period. The food habituation was designed with a clause to add an extra day if the reward options were not consumed by both calves in the pen after the three days, but this scenario did not occur for the enrolled calves.

#### 2.3.2. Training

Three days prior to the experiment’s start date, all the calves underwent a training and habituation process using a simplified version of the puzzle box with no intentionally challenging elements. The purpose of the training was to get the calves accustomed to entering the 0.8m corridor behind the pen using the first door (D1), approach the enrichment area to interact, and exit the area back into the pen via the second door (D2). Moreover, we wanted the calves to know where the food reward was located so that they could be motivated and comfortable to seek it out once the puzzle elements were present. A layout of the pen and experimental area can be found in Figure 3.

**Figure 3.**
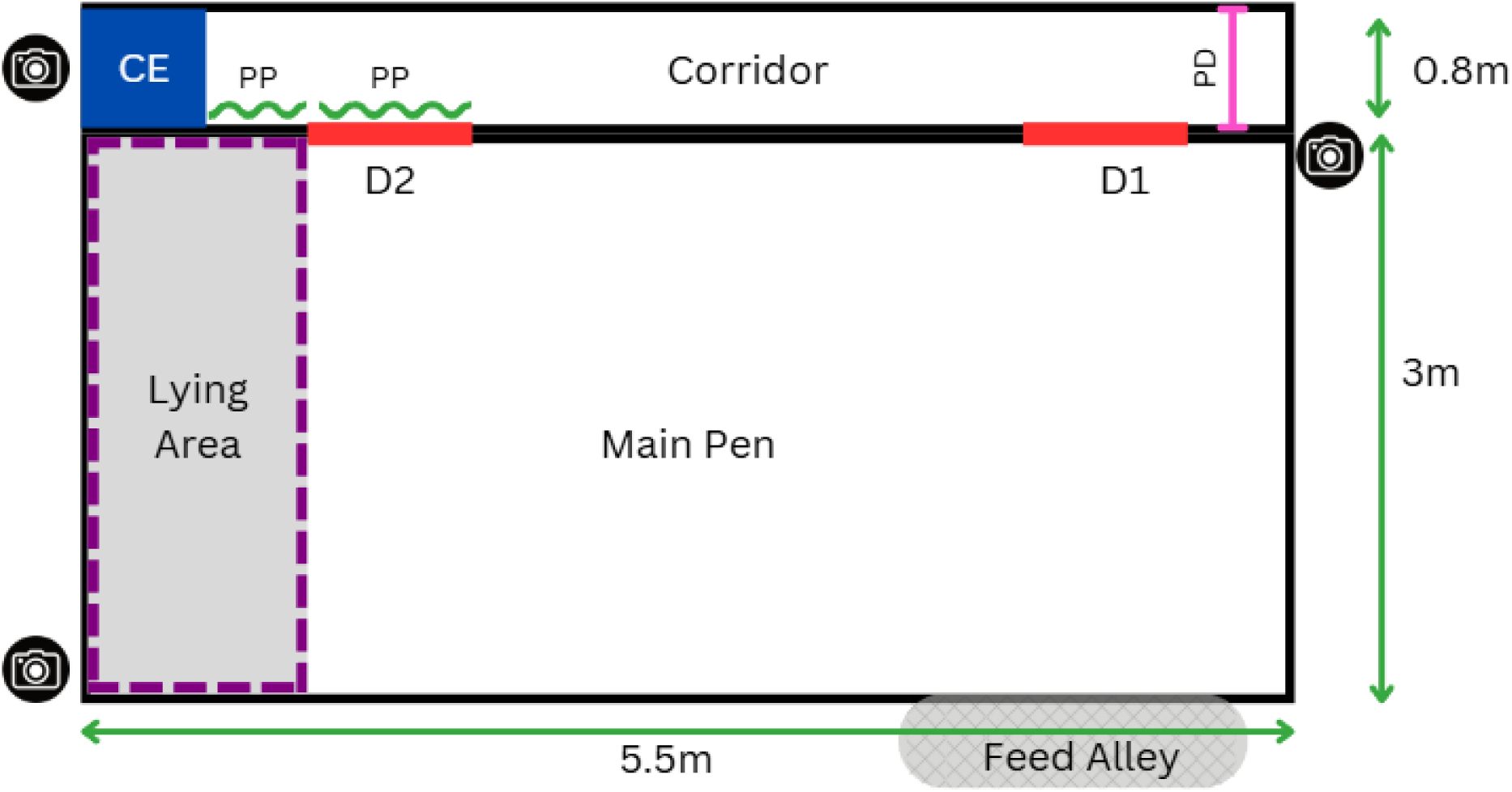
Illustration of calf pen and the experimental area. Calves enter the corridor via Door 1 (D1), move down the corridor to the cognitive enrichment area (CE), then exit via Door 2 (D2) to rejoin the main pen. Coroplast privacy panels (PP: 122 x 75cm; 122 x 47cm), hide the participating calf from the main pen and are located on D2 and in front of CE. The panel door (PD: 76cm x 121cm) is located right behind D1to act as a physical barrier from the rest of the corridor.

For the three training days, the cognitive enrichment (CE) area was comprised of a 43cm x 41cm wooden box with an open front that was mounted on a modified wooden fence post (see Figure 4). The box was mounted at shoulder height of the calves (approximately 1.23m) and the whole apparatus was fixed in place using two cut pipes secured to the pen gate. The calves were unable to see or pass through the CE area because the apparatus was fitted with white coroplast panels to close all visible gaps. For day 1, the box remained open faced but for day 2 and 3, we equipped the front with a transparent Plexi-glass panel that had a rectangular hole cut into the lower half large enough to accommodate a calf’s muzzle (approximately 12 x 14cm). A combination of two out of the three food reward options (cabbage x banana; cabbage x watermelon; watermelon x banana) was randomly selected for each day and approximately 2Tbs of the cut and processed mixture was placed inside the box. Prior to the training start, any remaining feed in the feed alley was removed for the duration of the training trial to promote motivation for the food reward. The feed was then promptly returned after the trial was over.

**Figure 4.**
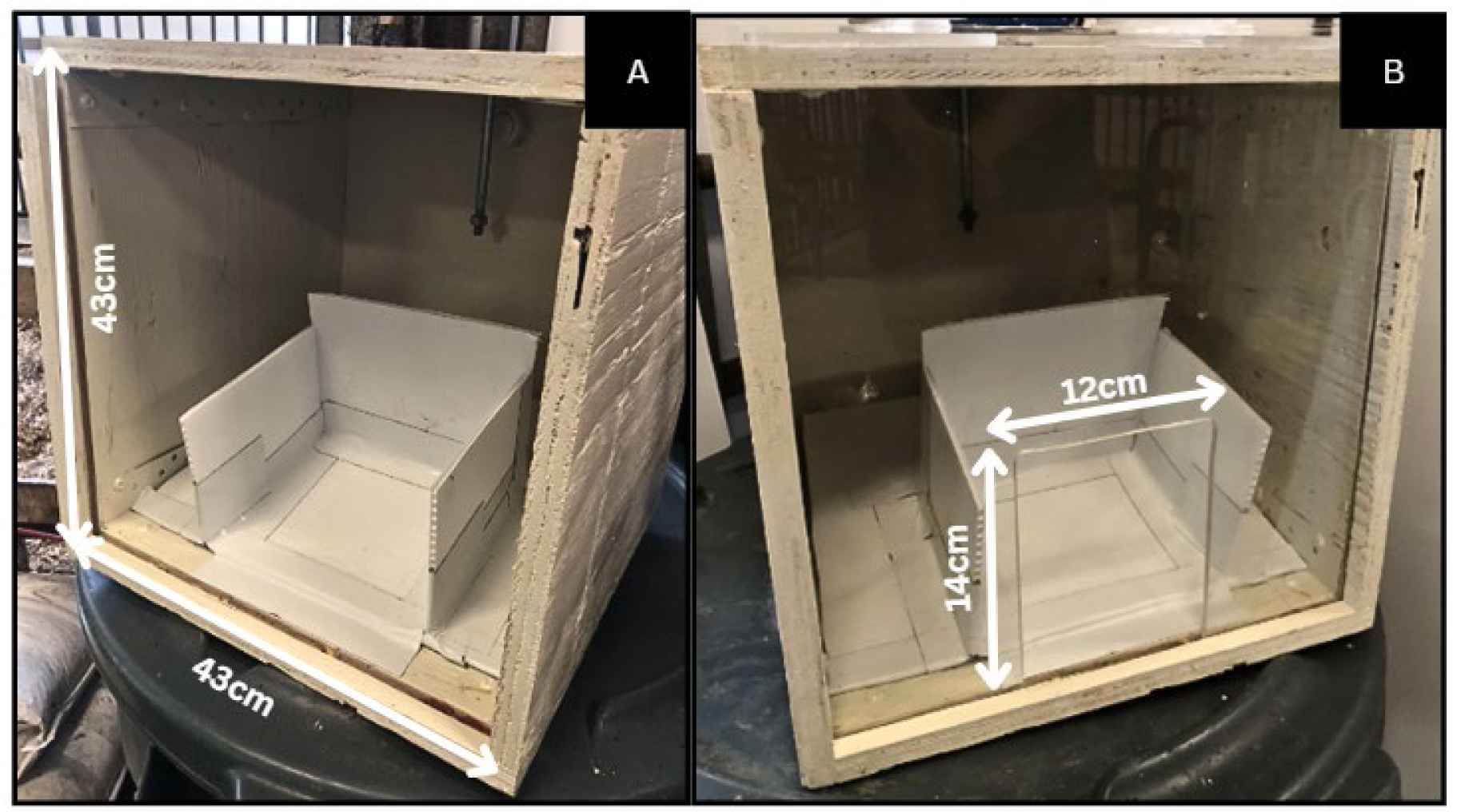
Training box for day 1 (A) is an open face cube with no CE elements. The training box used for day 2 and 3 (B) is structurally identical to the ‘day 1’ box with the addition of a Plexi-glass panel fitted to the front with a hole cut out of the lower half to fit a calf’s muzzle.

The criteria for passing day 1 of training involved having each calf inside the pen enter the corridor (either voluntarily of human-led), approach the CE area and retrieve the reward from the box. Each calf had two attempts on day 1 and if an individual failed, she would receive further training within the same day. There was an overall maximum session limit of 20 minutes (or 10 min/calf) to reduce stress if the animal was not responding well to the training. However, this was not an issue we encountered across the five groups. The order of calves’ participation (entering the enrichment area), the environmental status (weather and physical state of the barn) and disruptions were recorded as live observations. Throughout the training process, three researchers were present: Person A, Person B and Person C. Person A was situated at the end of the corridor behind the panel door and ensured the calf returned to the pen after visiting the CE area. Person B remained inside the pen to moderate the individual CE area access and Person C was outside the pen taking observations and giving cues.

Person A was trained to use an escalatory 5-step exit method for guiding the calves out of the corridor once Door 2 was open. The exit method is described here because its data contributed to our analysis. The procedure was as follows and Person A would only progress through the steps if the calf did not exit the corridor: Step 1: For the first 30 seconds of Door 2 being opened, Person A would not engage the calf. Step 2: After 30 seconds, Person A would move in front of the PD and approach the calf while clapping and using vocal encouragement. Step 3: After another 30s, Person A would add gentle physical touch in the form of rubbing and/or tapping the calf’s rump. Step 4: After another 30s elapsing, Person A would gently but firmly push the calf towards the Door 2 exit. Step 5: If another 30s go by, person B would assist person A to get the calf back in the pen using a food lure or guiding with their hand.

For days 2 and 3, calves had one 20-minute trial in which they were able to voluntarily access the CE area individually. During this time, individual access was monitored and maintained by researchers. The calves were able to go as many times as they could during the time limit and the only restriction was that minimal requirement of each going at least once. If one of the calves did not voluntarily visit the CE area by the 10-minute mark, she was encouraged to go using verbal and light physical handling (rubbing or patting on the rump). The only change between the two days was the addition of coroplast privacy panels (see Figure 3) introduced on day 3, on Door 2 and the gate in front of the CE area, to create a visual barrier between the CE area and the pen. The privacy panels were withheld until now under the premise that the calves would experience minimal stress with the space being visually isolated, since they were previously habituated to the area. If both calves in the pen were able to successfully retrieve the food from the box three visits in a row over the course of the last two days, then they could proceed to the experimental testing stage. If one or neither did not meet the requirements, additional training would be invoked until the criteria was met. However, none of the calves in our cohort required additional training. Live observations for days 2 and 3 of training included frequency of participation, order of participation, behavioral observations (such as exploratory or fearful interactions) and environmental status. For the control group of our cohort, they continued with day 3 training set up for the following nine-day experimental period.

### 2.4. Experimental Phase

#### 2.4.1. Puzzle Boxes

The cognitive enrichment for this experiment consisted of three different puzzle box doors with varying solutions that were fitted onto the wooden box and mount used in training (see Figure 5). The puzzle boxes were designed to be solved using different oral manipulation actions that are part of the calves’ natural exploratory behaviors. The puzzle box front panels were made of transparent Plexi-glass and had colorful rope integrated into some sections of the door designs. The rope was chosen to be a textured element that the calves can use to more easily orally manipulate and interact with the puzzle door. The colors were chosen to be part of the calves’ visible spectrum (Phillips & Lomas, 2001) and act as an attractant to promote interaction.

**Figure 5.**
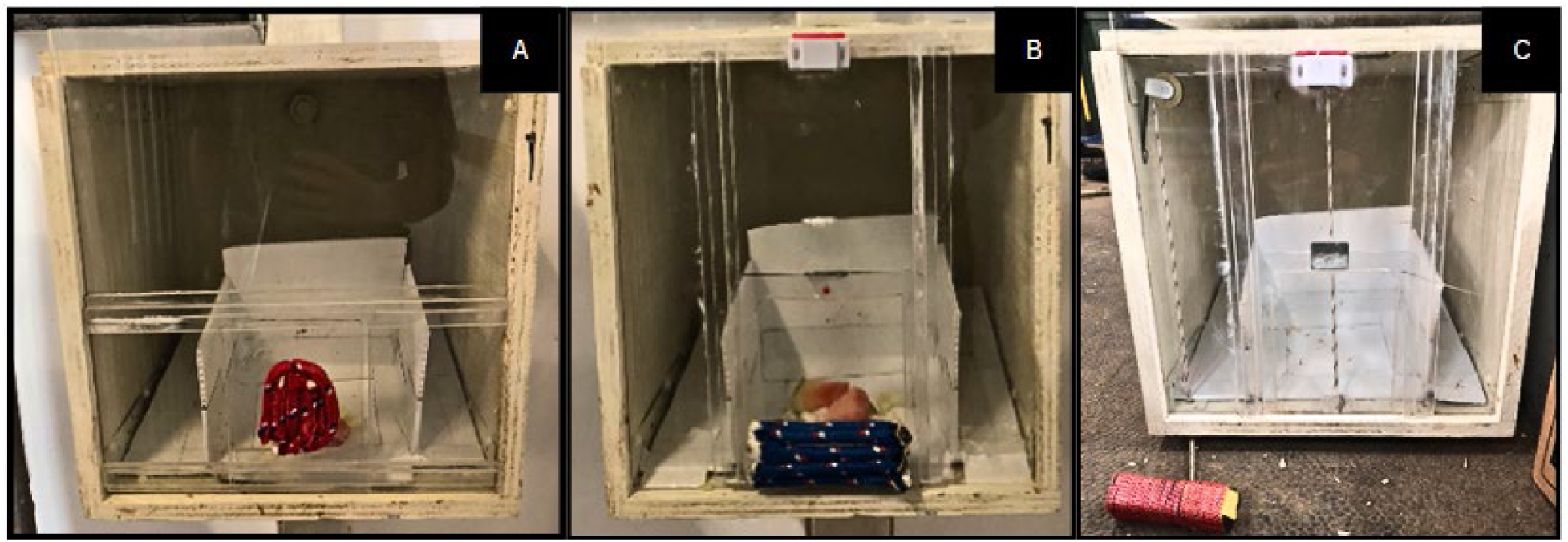
The three puzzle boxes: Slide box (A), where the door moves on the horizontal axis. Push box (B), where the door moves on the vertical axis. Pull box (C), where the door moves on the vertical axis when manipulating the multi-directional handle.

Firstly, the Slide box (see Figure 5 A) had a door fitted on a railing and could be moved on the horizontal axis using a plastic protrusion wrapped in red rope. In order to solve the box and access the food inside, the calf had to slide the door all the way to the left or right. Secondly, the Push box (see Figure 5 B) had a door that moved on the vertical axis and had a thicker rectangular ledge on the bottom that was outfitted with blue rope. The door could be opened by pushing the door up until it connected magnetically to the top of the box and would remain ajar unless it was pulled back down. Lastly, the Pull box (see Figure 5 C) door was completely smooth on the outside and was designed to be opened using an external handle. The door was attached to a pully system operated by a rectangular plastic block wrapped in red and white rope hanging off the end of a string. The door could be opened by pulling the block in any direction to create tension on the sting that in turn, pulled the door up. Similar to the Push box, the door had a magnet at the top of the box that kept the door open once it reached the appropriate height.

#### 2.4.2. Experimental Procedure

The experimental groups had the opportunity to experience all three boxes over the course of nine days. The order in which the puzzle boxes were presented to each group was partially randomized through a random number generator with a restriction on consecutive box repeats. The order was randomized because the puzzles were designed to be of equal difficulty level and not a ranked order. The frequency of box swapping during the nine-day experimental period varied depending on when the calves met the set criteria. The criteria made it so once a box design was solved (ie: opening the door) three times consecutively by at least one of the calves in a respective pen, the box design was changed to another. This had the potential to occur either over the course of two trials or one. We chose to go with three consecutive successes because that assumed that the solving ability was not a coincidence past this point. Moreover, we did not want a higher success rate threshold in order to keep the novelty of challenge and avoid the puzzles becoming a less cognitively stimulating operational enrichment. If a box was not solved according to the criteria after 6 trials (3 days), then the box design was changed to maintain the novelty.

The experimental procedure began each day at around the same time for standardization. The procedure was identical in format for both control and experimental groups save the type of box presented in the CE area. Calves participated in two 20-minute cognitive enrichment trials per day (18 trials total), both of which followed the same procedure for all 9 experimental days. The daily trials were separated by a 10-minute intermission during which calves were denied access to the CE area, and all personnel vacated the barn to minimize external influence. Each trial began once all cognitive enrichment elements were installed, the remaining food was moved away from the pen and the cameras were set to record. The CE set up followed the day 3 training set-up, except one of the puzzle boxes replaced the Plexi-panel box front for the experimental groups (see Section 3.3.4.1. for details on boxes). Three cameras were used to record the experimental trials: two GoPro HERO11 units mounted in opposite corners of the double pen for a wider view of the pen and corridor, and one AXIS M5074 PTZ network camera (Axis Communications AB, Lund, Sweden) positioned directly above the CE area for a closer view of the enrichment area (see Figure 3). The 20-minute timer would start once Door1 was fully open. The voluntary participation and one-at-a-time access system was identical to that of the training phase and all calves had at least one CE interaction per trial. The only time researchers interfered in the calf voluntary participation was if both calves tried to enter the corridor at the same time or if a calf needed to be encouraged to meet the minimal access requirement. If two calves tried to enter at the same time, Person B would use their best judgement to hold back the calf that is furthest away from entering the corridor.

Each time a calf entered the corridor to access the cognitive enrichment, she had a maximum of three minutes to solve the puzzle. This time limit was to maintain interest in the cognitive enrichment. The end of a calf’s visit was determined by opening Door 2 for the calf to return to the home pen under four scenarios. First, if the maximum three minutes of interaction was reached. Second, 30 seconds after the calf fully consumes the reward inside the box. Thirdly, if the calf has disengaged with the CE area for longer than 15 seconds (even when the puzzle was unsolved). Fourthly, if we observed a false visit, which was when two calves entered the corridor at the same time or a calf turned around in the corridor. With false visits, the 5-step exit method was skipped, and the visit was promptly terminated to avoid possible injury. Each time a box was solved or assisted solved (see below for details) during the trials, the reward inside the box would be replenished, and the puzzle would be reset before Door 1 would re-open for calf voluntary access.

In order to promote engagement with the puzzle boxes, a randomized assisted solve method was deployed for calves unable to solve the puzzle. If a calf consecutively failed to solve the puzzle and ceased to interact with the cognitive enrichment elements, we would open the box door at the 2-minute mark. This procedure was applied only after 2-3 consecutive failed attempts and randomly selected, to avoid discouraging puzzle interaction. The calf then had a maximum of one minute to approach and take the reward from the box. If she did not retrieve the reward, Door 2 would open and the calf would be returned to the pen. There was no endpoint for if calves continuously failed, thus calves would only be removed from the experiment under health/ welfare reasons. However, we do acknowledge that continuous failures carry the potential to frustrate and stress the animals. We tried to mitigate this outcome with the randomized ‘assisted solves’ that would occasionally provide the calf with food reward regardless of performance.

### 2.5. Measures

#### 2.5.1. Behavioral Measures

Live observations were taken to record the frequency and order of participation, in addition to the solving rate of the puzzle boxes as a pass/fail during the trials. The PTZ video files were used to record behavioral interactions the calves had with the cognitive enrichment and surrounding area, while the GoPro video files were used for recording calf behavioral measures within and around the enrichment area and corridor. The measures we took were the latency to approach the enrichment, the durations of visits, behaviors and calf exits (Door 2 open), the participation frequency and success rate (defined in Table 1, Table 2). The behaviors listed above were analyzed through video annotations completed by two blind observers using the software program Noldus Observer XT 14 (Noldus, 1991). The observers were trained by an ethology expert using a series of practice videos and a gold standard video inclusive of all behaviors outlined in the ethogram. Inter– and intra-observer reliability was checked within the software program every 10 videos using the gold standard, to ensure a minimum of 85% reliability.

**Table 1.** Type and description of behavioral measures taken during live observations and video analysis.

| Behavioral Measure | Description |
| --- | --- |
| **Latency to approach\*** | Time it takes for the focal calf to approach and contact the CE apparatus during each trial when given the opportunity to do so. Each latency starts when Door 1 is open and unobstructed, then stops when the calf initiates her first physical CE interaction. Specifically, touching with any body part or sniffing with muzzle at 1–10 cm from the object (Nawroth et al., 2017) |
| **Duration \*** | 1) Visit duration is from when the calf enters the corridor and initiates her first interaction with the CE to when she exits.<br>2) CE interaction duration is time spent interacting directly with the puzzle box.<br>3) Time it takes for a calf to exit the enrichment area once Door 2 is open. |
| **Participation frequency (Visits)** | The number of times each calf visits the CE area per trial. It is inclusive of voluntary and encouraged visits. Excludes false visits if no CE contact is made. |
| **Success rate\*** | Measured in ‘pass (1)’ or ‘fail (0)’.<br>Pass= The calf opens the puzzle box with no external aid during a visit.<br>Fail= The calf is unsuccessful at opening the puzzle box within a visit. |

\*= linked to visits which are the participation frequency within a trial.

**Table 2.** Ethogram of behavioral measures including type of behavior and description. This ethogram was used for observer training and analysis of calf behaviors in the CE area.

| Category | Behavior | Description |
| --- | --- | --- |
| **Interaction with CE** | Sniffing | Nose/muzzle is close to enrichment (within one muzzle length), while the calf inhales and exhales in a short repetitive manner (Manfre et al., 2024; Westerath et al., 2009). Slight movements of the nostrils may be visible, directed towards the CE apparatus. |
|  | Licking | Tongue is visible outside of mouth and is in contact with surface/CE materials at least once (Duve & Jensen, 2011). |
|  | Biting | Calf grabs at surface/CE material(s) using mouth/jaw. Movement of lips or teeth over the surface/ CE material(s) is visible. Includes ‘pull’ actions in which the calf bites a surface or object and pulls it in a direction, thereby displacing it from its original position. |
|  | Pushing | Calf uses her muzzle to press against a surface or object and exert a level of force against it (can vary from mild to aggressive). This can be in an upward, downward, left, right or forward action. |
|  | Head and neck rubbing + butting*neck | The calf contacts and rubs head or neck on a surface in a repetitive up-down or side to side manner (Ugwu et al., 2021; Whalin et al., 2022) or repeated bumps against the object (Westerhath et al., 2009). |
|  | Withdrawal (point event) | Calf abruptly pulls away with ears forward and eyes widened on the target. Often associated with a fear response and not to be confused with disengaging interaction (Whalin et al., 2022). |
|  | Eating | Calf must have some part of the muzzle or tongue inside the box. Eating can be defined as taking food into the mouth followed by moving jaw in a chewing motion and swallowing (Mac et al., 2023). If eating outside the box, code as a non-CE interaction. |
|  | Other | Any other behaviors not specified in the ethogram. |
| **No Interaction** | Standing/ walking | Calf is either standing on all four feet with minimal body displacement or directionally walking and not interacting physically with the environment. Includes self-grooming in which the calf is using her tongue to lick parts of her body or using her feet to scratch an itch. |
|  | Environmental interaction | Any type of interaction with the surrounding environment and anything that is not the puzzle box or associated elements. This includes the floor, the CE mount, Privacy panels, metal gate, researcher, walls and another calf. |
| **Extra** | In pen | Calf has put her two front feet into the home pen. If she backs back into the corridor (ie: back feet in the pen first), take the final time she puts her front feet into the pen as the code. |

Weather conditions were recorded daily to gather contextual information that may explain potential outliers and unexpected results. Prior to each trial, the temperature (°C) and wind speed (mph) inside the barn were recorded. Furthermore, any deviations from standard routine such as veterinary visits, off-schedule cleanings or feedings and loud external disturbances were also recorded.

## 3. STATISTICAL ANALYSIS

### 3.1. Raw Data Manipulation

The latencies to interact with the cognitive enrichment were analyzed to examine if they changed through time and if there was a relationship to success rate. Since success rate was initially recorded as “pass/fail”, we converted the data to binary 0=fail and 1=pass denotation. Latency times were expressed as seconds. Since the pair housed calves were provided with enrichment access at the same time, we had to subtract visit duration of one calf from the pen mate’s latency if overlap occurred. For example, if Calf 1’s first and second visit sandwiched Calf 2’s first visit, then we would subtract Calf 2’s visit-one duration from Calf 1’s visit-two latency. This is because during a calf visit, the CE area is unavailable to the other pen mate since the doors are closed. Therefore, we removed the latency time of when the access was barred for a more accurate representation of their voluntary participation. All durations were expressed in seconds. The duration proportions for two analyses were calculated directly in R and added to the data table. Firstly, the total group proportion of time spent in the CE area was calculated by adding the CE area total durations for both calves within a group and then divided that number by the total trial time which was then multiplied by 100 to get the proportion in percentage. This was done because calves within a group affected each other’s individual CE area durations and in turn, the total proportion of total trial time spent in the CE area. Secondly, we took proportions for direct interaction within the CE area. This was calculated by dividing the total direct interaction duration of calves (at the individual level) by the total CE area duration and multiplying by 100. This was done at the individual level because the other calf does not have an effect on the proportion of CE area time spent interacting directly with the puzzle boxes. For descriptive stats, we used the raw data from Door 2 exit durations categorized in 30 second time increments representative of the 5-step exit strategy to look for trends. For the behavioral data, we analyzed the frequency and durations of all the behavioral expressions outlined in the ethogram.

### 3.2. Data Analysis

#### 3.2.1. General Data Analysis

Descriptive stats for visualizing trends in data for durations were conducted through Excel (Microsoft Corporation, 2024). All statistical analysis was performed in R Studio version 4.4.1 (R Core Team, 2023). The following packages were used: *dplyr* (1.1.4; Wickham et al., 2023), *ggplot2* (3.5.1; Wickham, 2016), *lme4* (1.1-35.5; Bates et al., 2015), *car* (3.1-2; Fox & Weisberg, 2019), *lmerTest* (3.1-3; Kuznetsova et al., 2017), *emmeans* (1.10.4; Lenth, 2023), *nlme* (3.1-166; Pinheiro et al., 2023), *gridExtra* (2.3; Auguie, 2017), the *tidyverse* collection (2.0.0; Wickham et al., 2019) and the *conflicted* package (1.2.0; Wickham, 2021) which was used to manage function name conflicts. Packages specific to individual tests and analyses are detailed in the sections below.

The same modelling methodology was used for all variables. It started with fitting a model using the try function (R base package) to allow error recovery, starting with a simple model and then gradually adding factors (fixed, random), nested factors (fixed, random), factors interaction (fixed), and covariance structure for repeated measures, until models that failed to run and why were identified. Most model failures were due to factors being confounded and the model being too complex. Hence, models that best represented the experimental design and yielded no errors in the prior step were selected to continue. In a second step, because the data contained multiple measures from the same subject, residual independence was assessed with the autocorrelation function (ACF) in conjunction with the partial autocorrelation function (PACF). Based on the significance and the pattern of the correlation observed in the ACF and PACF graphs, the nature of the required covariance structure was defined or confirmed not necessary if no correlation was detected. All variables in the latency and duration analysis had no residual dependence hence did not benefit from the addition of a covariance structure in the model and were analyzed with a simpler model with fixed effects and a random intercept. As for the variables in the behavior analysis, residuals were not independent so an attempt to fit the best covariance structure to control for the dependency of residuals was conducted but failed to find a structure that would handle the residual dependency either because it was not a good fit (did not improve the model) or failed to run because of the unbalanced data. Other options were attempted to handle model misspecification such as robust SE, GEE, robustlmm, and addition of a random slope but none of the analysis worked likely due to the small sample size, and complexity of the model. Thus, the final model for the behavior analysis does not completely handle residual dependence and we acknowledge the potential for type 1 error inflation boosting inference.

Once the final model determined for all analysis types, the normality and variance homogeneity were assessed visually for all variables using Q-Q plots, and histograms. When normality was not met and/or the variance shown heteroscedasticity, the appropriate data transformations were applied (Table 3). Statistical significance of fixed effects was evaluated using Type-III ANOVA (Satterthwaite’s method) for denominator degrees of freedom. For post-hoc pairwise comparison, a Tukey, Bonferroni, Dunnett or Scheffe correction was applied based on the type of contrast. Finally, to investigate systematic trends of variables across trials, we applied polynomial contrasts and assessed the significance for linear, quadratic and cubic trends. All statistical results are reported using transformed data. Significance was declared at P ≤ 0.05, and tendencies were set between 0.1 and 0.05.

**Table 3.**
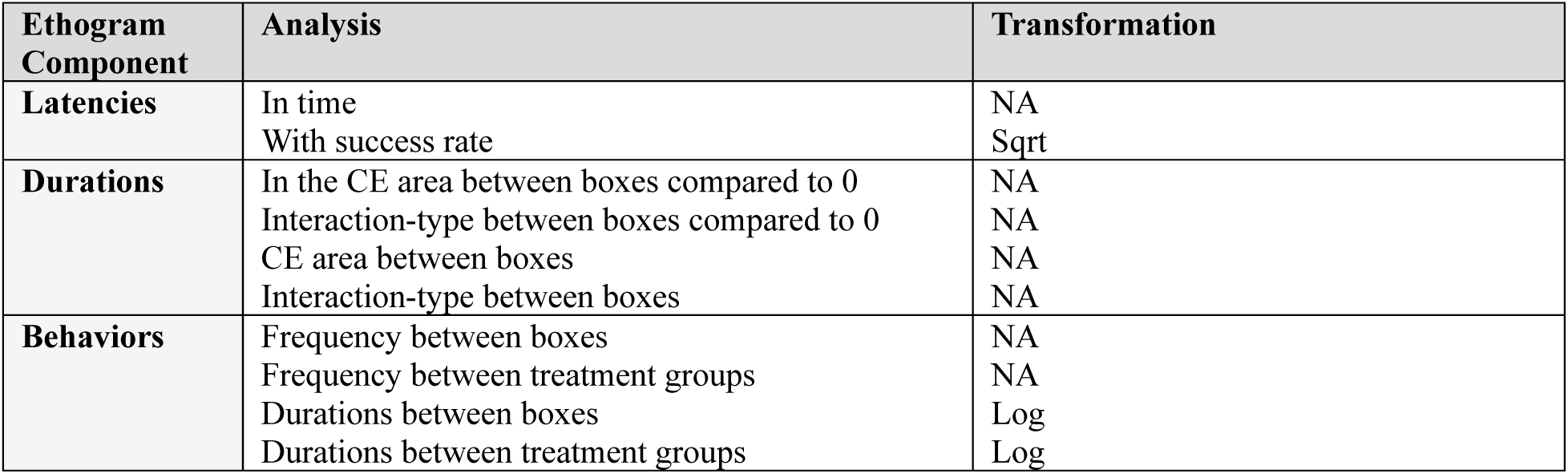
Data transformation to reach normality of residuals.

#### 3.2.2. Latency Analysis

The latencies of calves to access the CE area on the individual level, was done using a linear mixed-effects model with Trial, Success Rate (SR), and Group as fixed effects. The fixed effect of the interaction between Trial and Success Rate was included to assess whether the effect of Success Rate varied across trials. To account for the hierarchical structure of the data and repeated observations, we included random intercepts for Visits nested in Trial (experimental setup) and Calf nested in Group. Post-hoc pairwise comparisons were conducted using Scheffe adjustment for analysis. The final model was specified as:

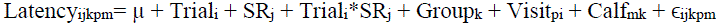

Where *i*= trial number (1-18), *j*= Success Rate level (0,1) *p*= visit (1-8), *m=*calf (1-8) and *k*= group (1-4 treatment groups). *µ*= the intercept and ε is the residual error term. Lastly, in order to further investigate systematic trends across trials, we applied polynomial contrasts (linear, quadratic, cubic) to the Trial factor by Calf, and by Success Rate.

#### 3.2.3. Durations Analysis

The analysis of durations was done in four parts. Post-hoc pairwise comparisons were conducted using Bonferroni adjustment for analysis.

##### 3.2.3.1. Is The Total Time Spent in The CE Area Different From 0 According to Box Type

The aim of the first analysis was to determine whether calves voluntarily spent significant time in the cognitive enrichment (CE) area, across trials and with the different box types, compared to 0. To address this, we analyzed the total duration of CE area interactions for each group (CEareaTdurs_sum) using a linear mixed-effects model. Trial and Box were included as fixed effects, and Group was included as a random effect to account for variability between experimental groups. Polynomial contrasts were applied to examine trends in CE area durations across trials. The model was specified as follows:

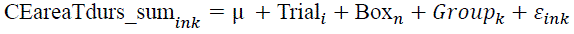

Where *i*= trial number (1-18), *k*= group (1-4 treatment groups) and *n* = box (slide, push or pull). *µ*= the intercept and ε is the residual error term. For trend visualization, we applied polynomial contrasts to observe the durations in the CE area in time (by trial).

##### 3.2.3.2. Effects of Trial and Box type on The Proportion of Time Spent in The CE Area

For the second analysis, we investigated if the proportion of time spent in the CE area (Perc_CEarea_sum) at the group level differed significantly based on box type, using a linear mixed model. Fixed effects included Trial and Box, while an intercept for Group was included for variation between experimental groups. The model was:

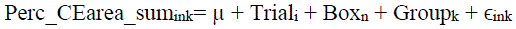

Where *i*= trial number (1-18), *k*= group (1-4 treatment groups) and *n* = box (slide, push or pull). *µ*= the intercept and ε is the residual error term.

##### 3.2.3.3. Interaction Type within the CE Area

The third analysis on the durations was conducted to understand how calves were spending their time in the CE area. Specifically, we looked into the interaction type which was split into either interacting directly with the enrichment or not interacting (environmental interaction or standing/walking). The durations of interaction type were assessed using a linear mixed-effects model with fixed effects Trial, Group, Interaction type (IntType), Box, and the IntType*Box interaction. The random effect was Calf. The model was specified as:

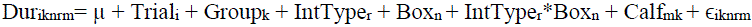

Where *i*= trial number (1-18), *k*= group (1-4 treatment groups), *n*= box (slide, push or pull), *r*= Interaction type (Interacting with the CE directly or not) and *m=*calf (1-8). *µ*= the intercept and ε is the residual error term. In addition to the mixed model, polynomial contrasts were applied to the trial factor for both interaction types.

##### 3.2.3.4. Effects of Trial, Group and Box Type on The Proportion of Time Spent Interacting with the CE

For the fourth and final durations analysis, we used a linear mixed effects model to assess the percentage of time spent directly interacting with the cognitive enrichment in time with fixed effects of Group, Trial, and Box. The random effect was Calf. The model was as follows:

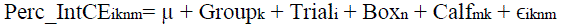

Where *i*= trial number (1-18), *k*= group (1-4 treatment groups), *n* = box (slide, push or pull) and *m*=calf (1-8). *µ*= the intercept and ε is the residual error term. We also applied polynomial contrasts to investigate any possible trends for interaction with the enrichment across trials.

#### 3.2.4. Behavioral Analysis

We separated the behavioral analysis into four main parts. The first two analyses looked at behavioral frequencies with and without accounting for treatment-factor. The other two analyses were on total durations of behaviors with and without accounting for treatment-factor. We chose to use both frequencies and durations because we were interested in the magnitude of behaviors expressed in addition to how long they performed each behavior. For both frequencies and durations, we did post-hoc pairwise comparisons with Tukey adjustment when excluding treatment-factor and Dunnett adjustment when including treatment-factor. We are initially excluding control to look at the differences specifically between puzzle boxes and then including control when aiming to look at the difference between treatments per behaviors. We decided to compare the treatment groups in the behavioral expressions analysis to note if the cognitive enrichment reveals differences in results from the control group.

##### 3.2.4.1. Behavioral Frequencies – Excluding Treatment

The model used for behavioral frequencies when excluding treatment from the model had fixed effects of Trial, Behavior, Box and Behavior*Box. The random effect was Calf. The model was as follows:

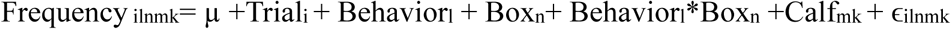

Where *i*= trial number (1-18), *k*= group (1-4 treatment groups), *n* = box (slide, push or pull), *m*=calf (1-8), and *l* = eight of the behavioral expressions (Sniffing, licking, biting, head/neck rubbing, environmental interaction, standing/walking, pushing and eating). *µ*= the intercept and ε is the residual error term.

##### 3.2.4.2. Behavioral Frequencies – Including Treatment

The model used for behavioral frequencies when including the treatment column had fixed effects of Trial, Behavior, Treatment, Behavior*Treatment and a random effect of Calf. The model was as follows:

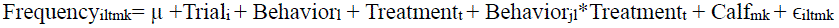

Where *i*= trial number (1-18), *l* = eight of the behavioral expressions (Sniffing, licking, biting, head/neck rubbing, environmental interaction, standing/walking, pushing and eating), *m*=calf (1-8), *k*= group (1-4 treatment groups) and *t*= the treatment group consisting of either the control box or the puzzle treatment boxes. *µ*= the intercept and ε is the residual error term.

##### 3.2.4.3. Behavioral Durations – Excluding Treatment

The model used for behavioral durations when excluding the treatment column had fixed effects of Trial, Behavior, Box and the interaction between Behavior*Box. The fixed effect was Calf. The model was specified as:

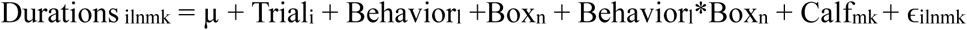

Where *i*= trial number (1-18), *k*= group (1-4 treatment groups), *n* = box (slide, push or pull), *m*=calf (1-8), and *l*= all nine of the behavioral expressions (Sniffing, licking, biting, head/neck rubbing, environmental interaction, standing/walking, pushing, eating and withdrawal). *µ*= the intercept and ε is the residual error term.

##### 3.2.4.4. Behavioral Durations – Including Treatment

And finally, the model used for behavioral durations when including the treatment column had fixed effects of Trial, behavior, Treatment, Behavior*Treatment and a random effect of Calf. The model was:

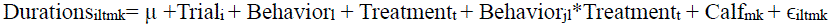

Where *i*= trial number (1-18), *l* = all nine of the behavioral expressions (Sniffing, licking, biting, head/neck rubbing, environmental interaction, standing/walking, pushing, eating and withdrawal), *m*=calf (1-8), *k*= group (1-4 treatment groups) and *t*= the treatment group consisting of either the control box or the puzzle treatment boxes. *µ*= the intercept and ε is the residual error term.

## 4. RESULTS

### 4.1. Latency to Interact

The latencies to interact with the cognitive enrichment for the treatment groups were assessed according to trial, group and success rate while accounting for variation among calves and visits. There was an initial significant effect on the interaction between trial and success rate (*F*= 1.67, *p* = 0.047). However, the post-hoc comparison revealed that none of the simple effects between success rate and trial for the latency remained significant after adjusting for multiple comparisons (all *p*>0.05). Estimated marginal means indicated that latency inconsistently fluctuated in time with success rate. Overall, the shortest latency recorded for a trip to the CE area was 1.80s while the maximum latency recorded was 708s (see Figure 6 for latency distribution). Across all trials, the mean latency was 75.7 ± 47.0s.

**Figure 6.**
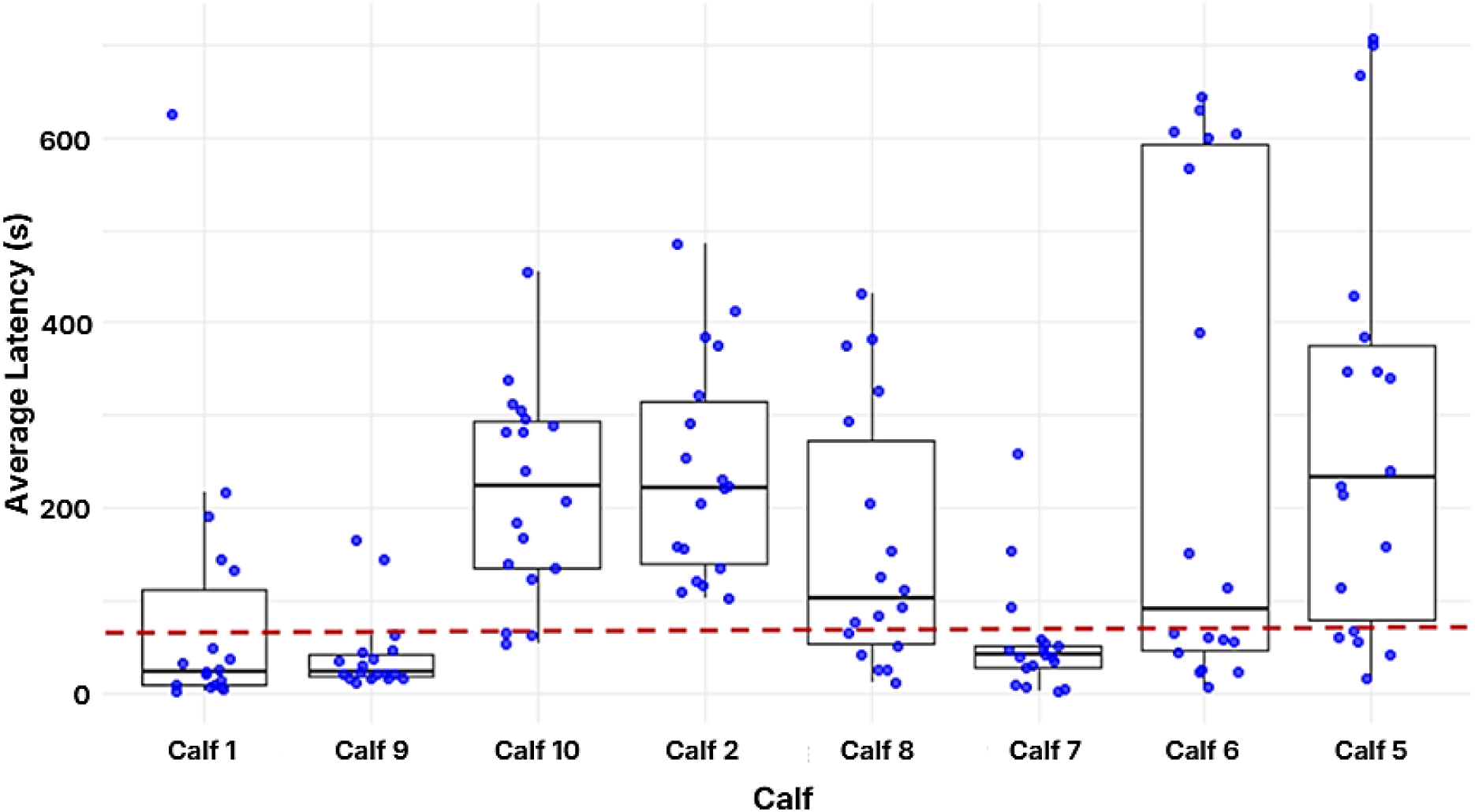
Boxplots showing the distribution of average latency (s) to interact with the cognitive enrichment (CE) by calf across 18 trials. Each box represents the interquartile range (25th–75th percentile) and the line inside the box indicates the median. Dots represent individual trial averages. The shortest latency recorded was 1.80s by Calf 7, and the longest was 708s by Calf 5. Across all calves and trials, the mean latency (red line) was 75.7 ± 47.0s.

To further our understanding of the calves’ motivations, we descriptively assessed the raw data latencies of encouraged visits to those where calves voluntarily participated (see Figure 7). Calves voluntarily participated for 87% of the total visits across trials with an average latency of 21.0 ± 5.20s. On the other hand, calves were encouraged for 13% of the total visits across trials with an average latency of 362 ± 26.0s. The average latency when succeeding was 72.1 ± 23.7s and for failing it was 78.5 ± 26.1s.

**Figure 7.**
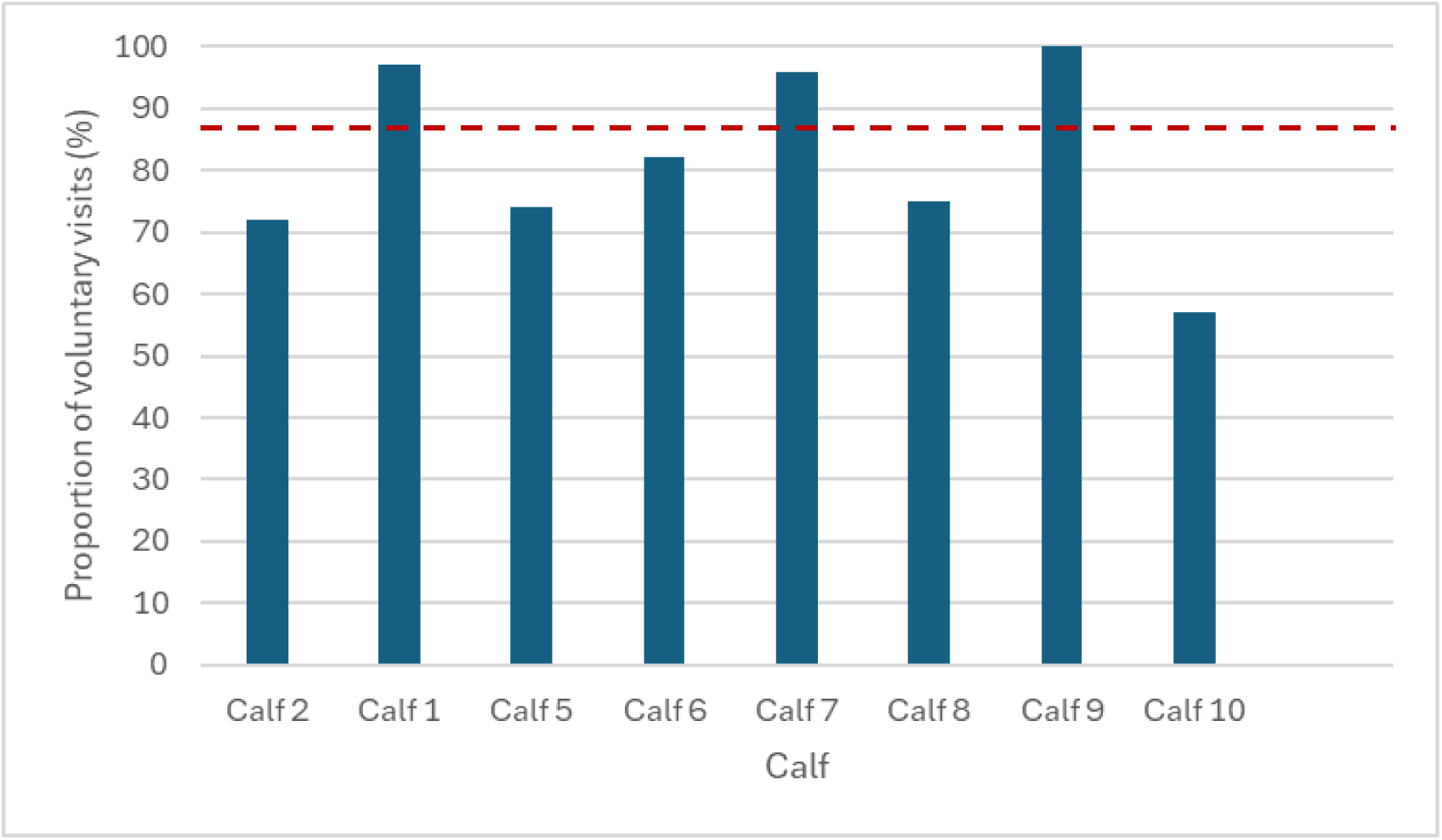
The proportion of voluntary visits to the cognitive enrichment area per calf averaged across all visits. The results demonstrated that calves voluntarily participated for an average of 87% of the total visits across trials (red line).

To examine trends in latency across trials, a polynomial contrast was conducted. The analysis revealed a significant cubic trend (*t*= 3.02, *p* = 0.003), whereas the linear and quadratic trends were not significant (all *p* > 0.10). These results suggest that latency did not change monotonically across trials but followed a nonlinear pattern with multiple inflection points, consistent with complex variable latency responses of calves throughout testing. A second polynomial contrast was performed on latencies by success rate to visualize if there were significant trends based on if the calves succeeded (1) or failed (0) the puzzle box. Results demonstrated that succeeding at the puzzle box had a positive linear trend of latencies across trials (*t*= –1.99, *p*= 0.05) and a significant cubic trend (*t*= –2.14, *p*= 0.035). On the other hand, failing the puzzle box gave the latencies across trial a significant cubic trend (*t*= 2.70, *p*= 0.008). These results indicate no clear differentiation in latency trend between a success rate of 0 (fail) and success rate of 1 (success) across trials. Thus, latency tended to fluctuate across trials independently of success rate.

### 4.2. Durations Analysis

#### 4.2.1. Is The Total Time Spent in The CE Area Different From 0 According to Box Type

Our first analysis assessed whether calves spent a significant amount of time in the cognitive enrichment area for each Box Type (push, pull, slide) across all Trials (1–18). On average, calves at the group level spent 65% (870 ± 21.0s) of the total trial time (1348s) utilizing the cognitive enrichment area, compared to remaining within the pen (35%; 478 ± 21.0s).

The linear mixed model revealed that compared to 0, calves spent a significant amount of time in the CE area for all the puzzle boxes (Slide: 748 ± 57.3s, *t*= 13.05, *p* <0.0001; Push: 894 ± 51.6s, *t*= 17.3, *p*<0.0001; Pull: 913 ± 47.1s, *t*=19.4, *p*<0.0001), while each trial was significantly different from 0 (range from 663 ± 88.5s to 1017 ± 87.6s; *t*= 8.13 to 11.61; All *p<0.0001*). These results suggest that engagement with the cognitive enrichment area fluctuated over repeated exposures. For a summary of the duration results, see Table 4.

**Table 4.**
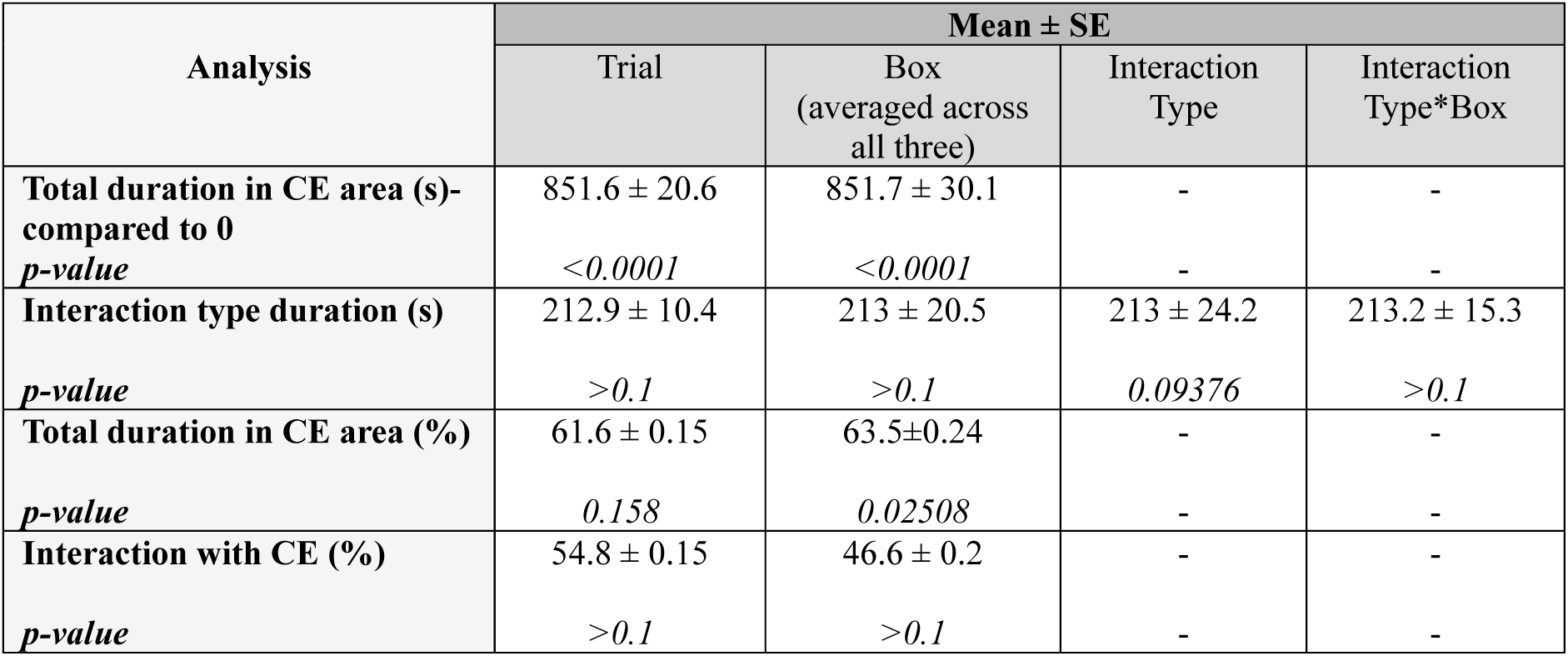
Summary of results reporting the mean ± SE for each duration analysis performed with the effect of each variable tested (Trial, Box, Interaction Type and Interaction Type* Box) plus the associated p-values.

| Analysis | Mean ± SE | | | |
| --- | --- | --- | --- | --- |
|  | Trial | Box<br>(averaged across<br>all three) | Interaction<br>Type | Interaction<br>Type*Box |
| **Total duration in CE area (s)-**<br>**compared to 0** | 851.6 ± 20.6 | 851.7 ± 30.1 | - | - |
| ***p-value*** | *<0.0001* | *<0.0001* | - | - |
| **Interaction type duration (s)** | 212.9 ± 10.4 | 213 ± 20.5 | 213 ± 24.2 | 213.2 ± 15.3 |
| ***p-value*** | *>0.1* | *>0.1* | *0.09376* | *>0.1* |
| **Total duration in CE area (%)** | 61.6 ± 0.15 | 63.5±0.24 | - | - |
| ***p-value*** | *0.158* | *0.02508* | - | - |
| **Interaction with CE (%)** | 54.8 ± 0.15 | 46.6 ± 0.2 | - | - |
| ***p-value*** | *>0.1* | *>0.1* | - | - |

#### 4.2.2. Effects of Trial and Box type on The Proportion of Time Spent in The CE Area

The effect of trial on the proportion of time spent in the CE area was not significant (*F* = 1.44, *p* = 0.158), indicating that time in the CE area remained consistent over time. In contrast, box type significantly affected the proportion of time spent in the CE area (*F*= 3.98, *p* = 0.025). Pairwise comparisons showed that calves spent significantly more time with the pull box (68.5 ± 3.8%) than with the slide box (58.6 ± 4.4%; t = 2.73, *p* = 0.027), whereas differences between the pull and push boxes (68.5 ± 3.8% vs 63.3 ± 4.1%; *t*= 1.6, *p* = 0.35) and between the push and slide boxes (63.3 ± 4.1% vs 58.6 ± 4.4%; *t*= 1.94, *p* = 0.71) were not significant.

#### 4.2.3. Interaction Type within the CE Area

Then, we compared the duration calves spent interacting directly with the puzzle boxes versus the surrounding environment within the CE area, across trials and box types. The model revealed no significant main effects of Trial (*F*= 0.61, *p* = 0.89), or Box type (*F*= 2.13, *p* = 0.12). The difference in duration between interaction types (direct interaction vs. non-direct interaction with the puzzle boxes), had a tendency towards being significant (*F*= 2.83, *p* = 0.094), with a mean comparison suggesting that calves spent slightly more time interacting with other elements of the environment (225 ± 34.2s) compared to directly with the enrichment (201 ± 34.2s). No significant interaction was found between interaction type and box type (*F*= 1.52, *p* = 0.22).

To explore trends in overall engagement with the CE area a polynomial contrast was conducted. No significant linear (*t*= 0.68, *p* = 0.50) or quadratic (*t*= 0.65, *p* = 0.51) trends were observed. A tendency for a cubic trend (*t*= –1.74, *p* = 0.083) suggested minor fluctuations in engagement over time. Overall, the results indicate that calves maintained relatively consistent durations of engagement within the CE area across repeated exposures, with no clear directional or cyclical pattern over time.

To further investigate the possible trend between interaction type and trial, a polynomial contrast was used on the predicted mean durations of calves by interaction type (directly interacting with the puzzle boxes and not interacting directly with the boxes). For direct puzzle box interactions, there was a tendency towards a cubic trend (*t*= –1.80, *p* = 0.073), while all other effects were nonsignificant (*p* > 0.1). This suggested that direct interaction with the puzzle boxes varied, but without a consistent directional change over time. Secondly, durations spent interacting with the surrounding environment showed no meaningful trend patterns (*p* > 0.05 for all contrasts).

#### 4.2.4. Effects of Trial, Group and Box Type on The Proportion Time Spent Interacting with The CE

No significant effects of Trial (F = 1.39, p = 0.15), or Box Type (F = 1.35, p = 0.26) were detected on the proportion of time spent interacting with the puzzle boxes. Estimated marginal means indicated that calves interacted directly with the boxes for an average of 43 ± 7.9% of the time spent in the enrichment area.

To further our understanding, a polynomial contrast was conducted to examine changes in the proportion of time calves spent interacting with the puzzle boxes across repeated trials, averaged over groups and box types. No significant linear (*t*= 1.27, *p* = 0.21) or cubic (*t*= –0.45, *p* = 0.65) trends were detected. However, a significant quadratic trend was observed (*t*= 2.20, *p* = 0.030), indicating that interaction with the puzzle boxes increased during the initial trials, peaked mid-series, and decreased slightly in later trials.

#### 4.2.5. Reluctance To Leave the CE Area

To further investigate the motivation of calves to remain in the enrichment area, we summarized the raw data durations of Door 2 being open (see Figure 8). We used this data to visualize the proportions of how long it took calves to exit the CE area across visits and trials. Results showed that 50% of the time (175 times), calves needed maximum encouragement and remained in the CE area over 91s past Door 2 being opened (step 4-5: gently pushed out or needed extra help from another person). Furthermore, calves remained in the CE area over 30s, 95% of the time (Steps 2-5; 329 times), meaning the researchers needed to intervene to get the calves to re-enter the pen.

**Figure 8.**
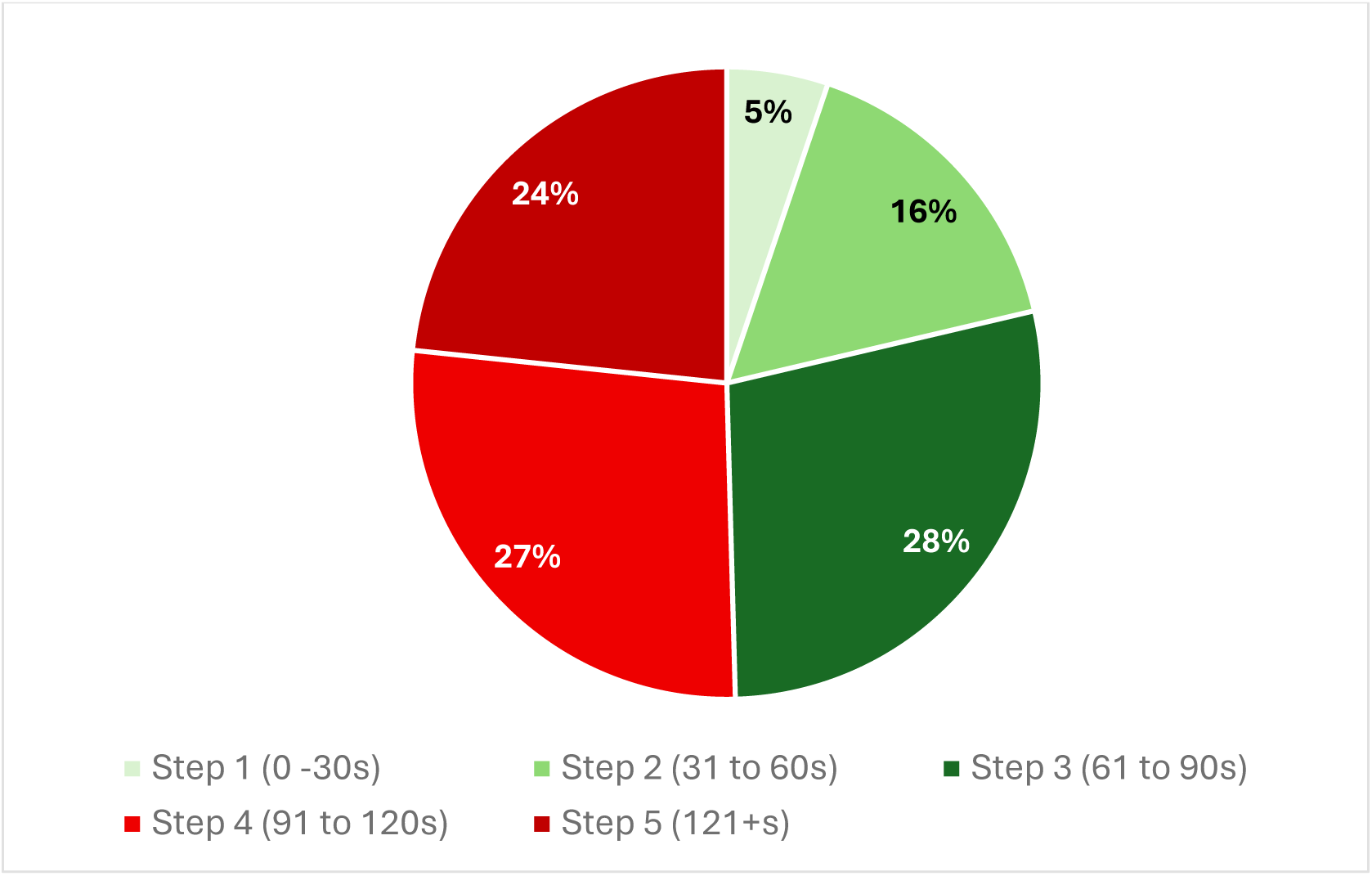
Proportion of how long it took the experimental calves to exit the CE area and re-enter the pen once Door 2 was opened via the 5-step handling method. The green to red color transition is to represent the increasing intensity of each handling step as they progress from 1-5.

### 4.3. Behavioral Analysis

#### 4.3.1. The Effect of Treatment on Behavioral Expression

A significant Behavior frequency*Treatment interaction (*F*= 7.72, *p* < 0.001) was found, indicating that the type of behavior expressed varied depending on if the calves received puzzle boxes or if they got the control box. However, the post-hoc comparison revealed that none of the simple effects between behavioral frequencies and treatment type were significant after adjusting for multiple comparisons (all *p*>0.05). See supplemental material Table S1. for detailed results of the treatment pairwise comparison. Estimated marginal means demonstrated that overall, sniffing (Control: 14.13 ± 2.44; Puzzle boxes: 17.10 ± 1.21) and environmental interaction (Control: 12.92 ± 2.44; Puzzle boxes: 11.22 ± 1.21) frequencies were among the most frequently expressed behaviors across groups, whereas biting, pushing, and withdrawal occurred infrequently (<4%).

The interaction between behavior durations and treatment was found to be significant (F=11.7*, p*<0.0001). Post-hoc contrasts comparing treatment (puzzle boxes) and control groups within each behavior (averaged across trials) revealed significant treatment-related differences for some behaviors. Calves with the puzzle boxes performed biting (*estimate* = 1.26 ± 0.44, *t* = 2.86, *p* = 0.013), pushing (*estimate* = 1.33 ± 0.44, *t* = 3.01, *p* = 0.010) behaviors for a longer duration than controls, and a tendency for licking (*estimate* = 0.81 ± 0.44, *p* = 0.091). Eating tended to be a lower duration in the treatment group (*estimate* = −0.90 ± 0.44, *p* = 0.063) than control. No significant differences between treatments were detected for environmental interaction, head/neck rubbing, sniffing, or standing/walking (*p* > 0.15 for all), though visual differences can be observed. Overall, these results indicate that the cognitive enrichment treatment selectively influenced specific behavioral expressions, particularly the oral manipulation behaviors, while overall engagement remained consistent across trials. See supplemental material Table S2. for detailed results of the treatment pairwise comparison.

#### 4.3.2. Behavioral Expression with Effect of Puzzle Box Type

The results revealed a significant main effect of the Behavior frequency*Box Type interaction (*F*= 4.30, *p* < 0.001). The significant Behavior frequency*Box interaction demonstrated that the relative expression of behaviors differed between box types. Random effects indicated moderate variability among calves (SD = 2.92). Overall, all of the nine behaviors were expressed to varying degrees across all box types. Behaviors such as sniffing, licking, and environmental interaction were associated with higher engagement frequencies, whereas head/neck rubbing, withdrawal, and pushing were expressed less frequently. Specifically, sniffing was the most frequently expressed behavior across all boxes (Slide= 17.6 ± 1.48; Push= 16.7 ± 1.35; Pull= 17.2 ± 1.27), whereas withdrawal was the least frequently expressed (Slide= 0 ±1.48; Push= 0.2 ± 1.35; Pull= 0.3 ± 1.27). Post-hoc Tukey-adjusted comparisons confirmed that sniffing, licking, and environmental interaction differed significantly from most other behaviors (*p* < 0.001), representing the primary exploratory responses to the puzzle boxes and CE area.

The results revealed significant effects of Behavior duration*Box Type (F=6.37, *p*<0.001), demonstrating that the effect of box type on engagement duration depended on the specific behavior performed. Estimated marginal means showed that environmental interaction was associated with the longest engagement durations across all box types with an overall average of 153 ± 30s. Apart from environmental interaction, sniffing and eating were the two other behaviors performed for longer durations for all puzzle boxes. For example, sniffing was performed for an average of 52.9 ± 11s (pull box), 44.2 ± 10s (Push box) and 37.52 ± 10s (Slide box). For the distribution of means for all behaviors according to box type, pairwise comparisons indicated that there were some significant differences in behaviors between puzzle boxes. For instance, pushing occurred longer on the Push (*t*= –4.02; *p*=0.01) and Slide box (*t*= –5.358; *p*<0.0001) when compared directly to the Pull box. Contrarily, some behaviors did not differ significantly between puzzle boxes such as biting and sniffing (all *p*>0.9).

## 5. DISCUSSION

The aim of this study was to assess the effects of providing cognitive enrichment to dairy calves. Specifically, we wanted to explore their motivation to participate voluntarily in the enrichment tasks and investigate how they interacted with them when given the opportunity. In order to accomplish this, we took various behavior measures such as latency to interact with the CE, durations within the enrichment area and a broad range of behavioral expressions. Combined, these measures can provide insights into the animals’ motivations which in turn can indicate whether providing cognitive enrichment can satisfy behavioral motivations and support positive experiences and welfare. We chose to focus on the motivation of calves to interact because sustained motivation to perform natural or rewarding behaviors indicates that the animal is capable of experiencing positive states like pleasure. Thus, through the introduction of cognitive enrichment, we attempted to analyze how the calves valued this addition to their environment.

### 5.1. Latency Analysis

#### 5.1.1. Latency as a Measure of Motivation

Studying the latency to engage with cognitive enrichment can provide useful insight into a calf’s motivation to participate and how familiarity with the task over repeated exposures can influence their response. For instance, in several cognitive learning task studies, latency to approach an operant target was considered to be one of the most direct measures of motivation to participate (Galhardo et al., 2011; Meagher et al., 2020; Ratuski et al., 2021). In addition to motivation, voluntary participation can indicate that some aspects of the experience were rewarding (Meagher et al., 2020), which is in line with the outcomes of an appropriate enrichment. In the context of our experiment, calves had the opportunity to participate voluntarily as many times as they could within the limitations of trial time and the availability of the enrichment area due to potential competition with the pen mate. To recapitulate, if the calf did not visit the enrichment by the 10-minute mark of the trial, she was encouraged to go. The calves within our experiment showed high levels of voluntary participation (87%) indicating that their interactions were not forced but rather consistent with internal behavioral motivations. These results also align with the positive animal welfare concept that providing animals with choice and autonomy can improve their welfare by fulfilling natural motivations and allowing animals to choose what is intrinsically rewarding to them (Rault et al., 2025). Overall, our findings indicate that calves were highly motivated overall to access the enrichment area and had relatively low latencies that fluctuated though time.

Attracting and sustaining an animal’s interest is an important hallmark of a successful environmental enrichment (Jones et al., 1991), and a good indicator that the animal is benefiting in some way from utilizing the enrichment. For our study, the cubic trend observed in latency across trials suggests a dynamic change in how calves approached the enrichment over time. Since the puzzle boxes changed throughout the trials at different rates, the fluctuations in latency are consistent with processes of learning which includes temporary frustrations and reduced novelty that can affect motivation (Kuhne et al., 2013; Bremhorst et al., 2019). Building on learning frustration, cattle that do not receive the anticipated stimulus when participating in a task that was previously rewarding, they can display reduced interest and interaction (Meagher et al., 2020). Therefore, it is possible that when the calves within our experiment received a new challenge, frustration may have occurred due to a box change. Specific to our cubic trend, early engagement could reflect curiosity and exploration, mid-phase latency increases could indicate lowered interest or learning frustration, and the later stabilization suggests the enrichment regained some sustained interest. These findings align with the idea that effective cognitive enrichment maintains animals’ interest by balancing predictability with cognitive challenge (Clark, 2017). The non-linear pattern observed suggests that calves continued to interact meaningfully with the enrichment over time, albeit with minor fluctuations to motivation, reflecting ongoing cognitive processing rather than habituation. It is important to note that while the calves in our experiment did not display habituation across the 18 trials, there is still the possibility of it occurring later on. Thus, in order to explore the topic of cognitive enrichment habituation further, we recommend a longer experimental time surpassing 18 trials or nine days.

#### 5.1.2. Influence of Success Rate on Motivation

In addition to the learning process, there is evidence that an enrichment with a foreseeable rewarding outcome creates anticipation through the association of an indicated stimulus to a significant event (Manteuffel et al., 2009). Essentially, the calves could be motivated to access the enrichment area due to the anticipation of receiving the high value food reward that comes with solving the box. We foresaw this as a possibility, which is why we assessed if the success rate had an influence on the latencies of calves. In the present study, latency did not differ significantly between success rate conditions, suggesting that the reinforcement reward offered did not substantially influence the calves’ willingness to approach and interact with the enrichment. Specifically, the overall experience of interacting with the enrichment could be positive, even for calves that failed the task. This is in line with the notion that cognitive engagement and learning, regardless of task outcome, can have benefits for animal welfare by providing an outlet for behavioral motivations (Meehan & Mench, 2007).

Furthermore, these findings may indicate that both conditions were sufficiently motivating on their own and that engagement was primarily driven by other factors like learning and mental stimulation, rather than external reinforcement. While some calves that failed their attempts at solving the puzzle boxes occasionally received some reward to reduce frustration and negative experience, this only occurred less than 33% of the failed visits. Due to its low frequency of occurrence and randomization, more often than not calves returned to the pen with no reward upon failing the puzzle. For the sake of acknowledgement, the anticipation of a food reward could have increased the overall attraction to the cognitive enrichment regardless of success rate, but it was not found to be directly related to the success rate.

However, that does not take away from the other rewarding elements that the calves may be gaining from interacting with the cognitive enrichment regardless of receiving a food reward. Similar results have been reported in other enrichment studies where novelty and individual exploratory tendencies outweighed reward-based differences in approach behavior. For instance, Rosenberger et al., (2020) found that goats were motivated to voluntarily participate in an operant task and were willing to work for a reward even in the presence of an identical, free reward. This suggests that the animals preferred to seek out a challenge and possibly found the task to be overall more rewarding and stimulating. They also reason that the animals may be choosing the operant task as a way to satisfy their exploratory needs and gain control over their environment. Additionally, De Rosa et al., (2003) supplied puzzle feeders to common marmosets and reported high levels of interaction with the devices even when not solving the puzzle. Specifically, the animals showed high levels of exploration and short latencies to approach the feeders, indicating levels of interest that were not previously observed directed towards their regular food dishes. While they mention the food reward had influence over the marmoset’s motivation, they also emphasize the exploratory satisfaction of interacting with a complex task. All this to say, the food reward used as part of our study could have contributed to the motivation of calves to access the cognitive enrichment, but since success rate did not significantly influence the latency, it is likely that other factors were responsible for the fluctuations in latency such as natural exploratory tendencies and learning. Particularly, we observed calves performing exploratory behaviors at different rates which could amount to individual differences in exploratory tendencies, resulting in natural fluctuations in latency.

#### 5.1.3. Summary

Since cognitive enrichment differs from other forms of environmental enrichment by intentionally targeting specific cognitive processes like memory and learning (Clark, 2017), there is promise that the added mental stimulation from solving a complex task could provide longer lasting engagement. With our experiment, we found dynamic interest in the enrichment over time, combined with proportionally larger voluntary interactions as valuable indications that the puzzle boxes hold potential as an effective cognitive enrichment for calves. Overall, the lack of success rate effect but presence of a dynamic trend highlights the importance of repeated exposure and cognitive novelty in sustaining calves’ interest in enrichment. These results support the view that young cattle can adaptively modulate their engagement based on experience, highlighting their capacity for flexible learning within a cognitively stimulating environment. Though further work is needed for understanding long term interactions and the possible changes to latency past a nine-day experimental time, there is reason to believe that calves are motivated to access cognitive enrichment when given the choice.

### 5.2. Durations

Building on the results of latency to access the CE, durations were used as a complimentary metric of motivation for investigating sustained engagement of the CE area and puzzle boxes. Specifically, calves were motivated to access the CE through latencies and durations were used to highlight the strength of the motivations by investigating how time was spent within the trials and visits. The analyses of durations within the enrichment area, durations of direct, indirect and puzzle box interactions plus the reluctance to leave the corridor, provided key insights into how calves engaged with the cognitive enrichment across repeated exposures. Collectively, the analyses of durations provide support that the cognitive enrichment was effectively motivating in promoting sustained and voluntary engagement.

#### 5.2.1. Durations in the Cognitive Enrichment Area

Overall, calves spent a substantial proportion of their total trial time within the CE area, indicating a strong motivation to engage voluntarily with the space. This suggests that the enrichment area was perceived as both stimulating and rewarding (Miranda et al., 2023). Moreover, the polynomial contrast of total durations in the enrichment area demonstrated consistency of engagement across trials, which provided further support that the enrichment area maintained its motivational value. While there is evidence of minor fluctuations in engagement, these are theorized to reflect natural variation in daily motivation or transient influences such as individual differences, social dynamics, or prior trial experiences. In other words, individuals differ in their responses to environmental stimuli due to internal and external factors which naturally create minor fluctuations in responses to enrichment (Zocher et al., 2020). Importantly, the absence of a downward trend indicates that the attractiveness of the cognitive enrichment area did not wear off, even after multiple trials.

In other enrichment studies with calves, decreases in engagement have been linked to repetitive or non-challenging enrichment tasks (Strappini et al., 2021; Zang et al., 2022), whereas consistent engagement has been associated with tasks offering cognitive stimulation or variability (Clark et al., 2023). Contrary to our findings, some studies suggest that habituation to enrichment occurs relatively quickly upon introduction. For instance, Strappini et al., (2021) found that most of the enrichment items (brushes, ropes and cowhide – note that none of these were intended to be cognitive enrichment, but rather physical enrichment) introduced to calves, received the highest number of visits during the first day of the study then subsequently decreased over time with a notable difference on day 4 of exposure and daily mean durations of under 75s. Furthermore, Van Os et al., (2021), observed that dairy heifers spent significantly less time using an originally novel brush (sensory) enrichment by the second and sixth day of presentation. While the true reasoning is unknown, it is possible that simple/predictable enrichments often experience faster habituation and thus, future enrichment should include elements that are specifically designed to adequately mentally stimulate and sustain the interest of calves.

Additionally, in previous enrichment studies, sustained voluntary engagement has been used as an indicator of motivation and welfare-oriented enrichment (Lambert et al., 2016; Rault et al., 2020; Taylor et al., 2023). These studies mention that sustained engagement may still be subject to modification by habituation over time in both short– and long-term contexts, but consistent stability suggests that the enrichment maintains the animal’s interest and provides an acceptable level of satisfaction. In other words, engagement with enrichment may naturally fluctuate in time but its consistent use is a good indicator that the enrichment is motivating and rewarding. Since the calves in our experiment consistently chose to remain in the enrichment area for the majority of the trial duration (even after repeated exposures) and their total durations did not fluctuate significantly, it is probable that the enrichment retained its attractiveness to the calves and did not induce habituation over the nine days. Therefore, the results here suggest that the enrichment configuration and delivery provided sufficient cognitive and sensory stimulation to sustain interest across repeated exposures.

#### 5.2.2. Interaction Type within the CE Area

For a deeper dive into how the enrichment area was used, we analyzed the distribution of time between direct and non-direct puzzle box interactions. Comparing direct interactions with the boxes versus interactions with the surrounding environment revealed that calves divided their time relatively evenly between these two activities with a slight tendency towards non-direct interactions with the puzzle boxes. The relatively balanced distribution of time suggests that calves were not only focused on the puzzle boxes themselves but also appeared to treat the CE area as an enriched microenvironment that invited broader exploration. Since calves are highly exploratory animals (Kerr & Wood-Gush, 1987; Vieira et al., 2012), it is unsurprising that they expressed interest in the surrounding area. Thus, even if the cognitive enrichment is engaging, environmental exploration is natural and expected. While they had constant visual access to the corridor outside of the experiment, they were unable to frequent the area and did not usually have the additions of the privacy panels and CE mount to explore. While one can argue the calves should have been habituated to the area due to repeated exposure throughout the experiment, there is one contributing element that kept the area continuously attractive. The calves tended to eat messily, and some proportion of the food reward ended up on the floor. When the calves ate off the ground it counted towards non-direct interaction and while we cleaned the corridor between visits, there was high probability that the floor was still highly enticing to the calves from residual food smells, influencing their interaction distribution. However, even with the sensory attraction of the floor, the calves still spent a large proportion of time interacting directly with the box enrichment.

Lastly, the non-direct interactions were split between environmental interaction and standing/walking. This later behavior can be explained by considering the results of other enrichment studies. Research on patterns of enrichment use in other species like mink, marmosets and rodents, have demonstrated that animals often alternate between active manipulation and observation or exploratory pauses, which may align with information processing or spatial learning (Meagher & Mason, 2012; Decker et al., 2023). The exploratory pauses within the enrichment area (indirect interaction of standing/walking) could be aligned with this conclusion. Furthermore, such patterns are consistent with studies in which animals exposed to cognitive tasks exhibit alternating bursts of interaction and observation, indicative of exploratory learning rather than loss of interest (e.g., Hagen & Broom, 2004). In summary, the interaction distribution suggests that calves tend to balance manipulation of the cognitive enrichment with natural exploratory behavior and pauses in interaction, reflecting a multifaceted engagement strategy. Therefore, the calves’ pattern of alternating between direct and non-direct interaction likely reflects a healthy form of exploratory engagement rather than a loss of interest in the enrichment. However, to confirm this we would need to look at behavioral sequencing, which is the order of how behaviors are layered and the order they occur (Vicino et al., 2022). All in all, the calves may be motivated to investigate the area in addition to using the device and taking momentary pauses to process, making the overall experience positive regardless of interaction direction.

While no significant main effects of trial were observed for direct or non-direct interaction durations, the polynomial contrast for direct interaction revealed a significant quadratic trend. This indicates that engagement with the puzzle boxes increased initially, reached a peak mid-series, and slightly decreased during later trials. Such a pattern could suggest an underlying learning or optimization process. Calves may have taken longer interacting with the puzzle boxes early on while they were learning the solutions to the puzzles. By the end of the experiment, all calves had learned to solve the three box variations which meant they were more efficient at opening the puzzle boxes and thus had shorter overall direct interactions. In other words, calves may have initially explored the puzzle boxes to understand their operation, achieved mastery by mid-trials, and then slightly reduced manipulation frequency once the task became familiar. This pattern is in alignment with the fluctuations observed in latencies. Thus, as mentioned with latencies, it is important to note that the decline in durations was not consistent with habituation since interest in the cognitive enrichment still remained relatively high and constant from the beginning to the end of the experiment. This interpretation aligns with studies in young cattle and other species like pigs, goats and chickens, that suggest cognitive tasks often elicit peak engagement once animals reach task proficiency, after which interaction rates stabilize or slightly decline (Lecorps et al., 2022; Clark et al., 2023). The small decline in later trials does not imply disengagement but rather a shift in behavior from exploration to efficient task execution. The stability of engagement across trials also speaks to the enrichment’s repeatability. In many enrichment studies, initial novelty can drive strong responses that fade as animals habituate. However, the consistent engagement patterns here suggest that calves found the enrichment inherently rewarding beyond the novelty phase. This may reflect the cognitive challenge involved, as problem-solving opportunities have been shown to evoke positive states in animals (Mellor, 2015).

#### 5.2.3. Influence of Box Type on Interaction Durations in the CE area

Results demonstrated that calves spent a significant amount of time interacting with all three puzzle boxes to different degrees. When compared between each other, the pull box had the longest average durations across trials, followed by the push box, with the slide box having the shortest interaction times. These findings highlight that engagement duration is not uniform across enrichment designs, suggesting potential differences in either perceived task difficulty, reward accessibility, or interaction preference. Enrichment design characteristics such as mechanical feedback and operability are known to influence animal engagement (Wilson et al., 2002; van der Staay et al., 2017; Vicino et al., 2022). Specific to our designs, calves faced unique challenges with each box that could have directly influenced box interaction even though we assumed the difficulties were equivalent. For instance, calves could have spent longer time with the pull box because it required some patience and skill to get the right leverage to grasp the handle and pull the door open. Secondly, the push box door had a height distance it needed to reach in order to remain open. If the door was not raised far enough, it could have taken more time to open. Finally, the slide door may have had a quicker solve time since once the door was pushed to the side, it would remain there. However, a downfall of the slide door was that the door opening was relatively smaller than the other puzzle boxes and could have created some difficulties accessing the reward once opened. In summary, all three puzzle boxes had their unique elements that could have increased or decreased how much time was spent directly interacting with the puzzle boxes due to perceived task difficulty and reward accessibility.

The trends in how long calves interacted with the puzzle boxes could also reflect a combination of task preference and differences in capabilities. Studies in pigs and cattle have shown that animals are more likely to persist with tasks that they can successfully complete or that produce a predictable outcome (Held et al., 2001; Hagen & Broom, 2004; De Jonge et al., 2008). On one hand, longer durations could represent a preference for interacting with the box, but on the other hand it could represent a higher effort invested in solving the box, thus requiring more time to interact. It is difficult to allocate which perspective the preference was on when a shorter duration could simultaneously indicate a faster solve and lower stimulation (i.e. the task was too simple). From a welfare perspective, this result emphasizes the importance of enrichment design where tasks can provide a balance between challenge and success in order to sustain voluntary participation and minimize frustration. For a more complete understanding of how the puzzle boxes were utilized, we discuss the behavioral expression of calves towards each box in later sections. Future enrichment protocols should consider how the form and feedback of cognitive devices align with the animals’ physical and perceptual capacities (Zhang et al., 2022). Thus, while further work is required to draw concrete conclusions, our study provides insights into developing appropriately challenging cognitive enrichment for calves.

#### 5.2.4. Reluctance to Leave the Enrichment Area

Perhaps one of the most compelling behavioral observations supporting our understanding of the calves’ interests, was their reluctance to leave the CE area once a visit ended. Specifically, within nearly all visits, calves required encouragement to exit, with half of these requiring maximal prompting. Given that calves were trained to associate the opening of Door 2 with returning to the pen, this resistance likely indicates that the enrichment area was perceived as a rewarding space in which the calves tended to overstay past the maximum time allotted per visit. This is further supported by the calves proportionally spending the majority of their trial time within the CE area by choice. Reluctance to leave a voluntary engagement area has been interpreted in other species like ruminants, pigs and non-human primates, as an indicator of sustained motivation and positive affective states (Webb et al., 2019; Meagher et al., 2020). In the context of our study, the calves’ behavior may reflect both cognitive curiosity and an expression of control over their environment, as they were able to freely choose when and how to interact with the enrichment. Such voluntary engagement is increasingly recognized as an indicator of a positive welfare experience, representing both agency and natural expression (Špinka & Wemelsfelder, 2011; Rault et al., 2025). Therefore, the calves’ resistance to leave the CE area paired with the results of other durations during trial and visit times, suggests the cognitive enrichment provided a meaningful and engaging experience.

#### 5.2.5. Summary

In summary, the duration analyses indicate that weaned dairy calves were actively and consistently engaging with the cognitive enrichment environment across repeated exposures. The enrichment maintained its motivational value over time, with calves spending the majority of their available time within the CE area and showing strong reluctance to leave it. Differences between box types highlight the influence of enrichment design on engagement, while the observed quadratic pattern in direct interaction suggests a learning-related adaptation rather than a loss of interest. Together, these findings demonstrate that cognitive enrichment can promote sustained voluntary participation and rewarding engagement, supporting the use of cognitive challenges as effective welfare-oriented enrichment for young cattle, promoting exploration, learning, and positive interactions with their environment. When combined with latency measures, the duration data suggest that calves not only approached the enrichment quickly but also maintained consistent engagement, supporting the utility of cognitive enrichment.

### 5.3. Behaviors

Investigating the behavioral expression frequencies and total durations of the treatment (with puzzle boxes) and control groups revealed valuable insights into the calves’ methods of interaction with the puzzle boxes and enrichment area. Across all groups and treatment types, calves performed a wide range of natural behaviors to varying magnitudes. These behaviors included sniffing, biting, pushing, licking, eating, environmental interaction, standing/walking, head/neck rubbing and withdrawal. Employing a broad range of behaviors towards the enrichment area suggests that the calves were overall motivated to engage with the enrichment and the surrounding area. This is due to the reasoning that the intensity of behavioral interactions with items can reveal the significance to an animal’s key motivations (Van de Weerd and Day, 2009).

Although the control and cognitive enrichment treatments differed in box design such that the puzzle boxes included cognitively demanding elements while the control box did not, both treatments support the opportunity for natural behavioral expression to some degree. It is possible that both treatment groups can create enough environmental complexity that facilitates behavioral expression in young calves. Furthermore, these findings imply that even a relatively small intervention can be meaningful in providing calves with an outlet for behavioral expression. For instance, simple additions such as ropes, balls and brushes can attract a calf’s attention and provoke interaction (Zobel et al., 2017; Strappini et al., 2021; Pereira, 2025). That said, the added cognitive challenge of the puzzle elements yields additional benefits in behavioral diversity (Milgram et al., 2006; Clark, 2017). Specifically, the puzzle boxes for the treatment group exhibited significantly higher behavioral durations when compared to the control group. Thus, the puzzle boxes in rotation could be considered multifunctional by targeting and satisfying multiple behavioral motivations in calves such as sniffing, licking, biting, pushing, head/neck rubbing and eating.

From a welfare perspective, the ability to perform motivated behaviors are an important component of good animal welfare (Hughes & Duncan, 1988; Rault et al., 2025). For example, multiple studies find that provision of non-cognitive enrichment items such as social, sensory, physical and nutritional environmental additions can increase behavioral diversity and reduce indicators of poor welfare in calves like inactivity and frustration (Mandel et al., 2016; Zhang et al., 2022; Occhiuto et al., 2025). In our study, the increased behavioral expression durations in the cognitive enrichment group paired with the sustained engagement observed in latencies and durations suggests that these calves had, arguably, a stronger match between their behavioral drives and their environment through the learning tasks provided by the puzzle boxes. On the other hand, the lack of cognitive task on the control box could explain the shorter behavioral durations since it was comparatively, a more restricted enrichment with fewer stimulating elements. Nonetheless, our findings support the argument that diverse options and opportunities to express a wide range of behaviors is an important key to fulfilling a calf’s behavioral motivations.

In the context of our experiment, the voluntary interaction configuration of the cognitive enrichment promoted choice in interacting with puzzle challenges. Though our experimental design had some restrictions like the one at a time method and the maximum 3min limit in the CE area per run, calves still had some liberty to leave their home pen and access the CE area of their own accord. Importantly, this choice was not something they had before the introduction of the cognitive enrichment. Furthermore, calves were unrestricted in their methodology of solving the puzzle boxes and could exert different techniques to achieve the same solution. For example, the push box could be opened by pushing the door up with their nose, biting the ledge or licking the door upwards. Combined, this increased agency allowed the calves to interact in their own way, which supports their individual problem-solving skills and development (Clark, 2017). Moreover, Rault et al. (2025) suggests that these environmental additions can help develop not only emotional, cognitive and behavioral competences, but also physiological and immune competences crucial to health and longevity. Finally, supporting motivated behavioral engagement and allowing animals to be in control is increasingly becoming a staple to rearing healthy, productive animals (Špinka, 2019; Colditz, 2022; Englund & Cronin, 2023).

Behavioral expression frequencies, durations and diversity are also relevant to cognitive development. Since calves use exploration and manipulation of objects within their environment as a means of processing and learning about their surroundings (Whalin et al., 2021; Nikkhah & Alimirzaei, 2023), it is important to provide outlets for these behavioral motivations. Exploration and manipulation through behavioral expression are things we observed high levels of within our experiment. In other enrichment studies, calves often spend more time manipulating objects with their mouths which is likely related to the young age of the recently weaned calves that retain a strong motivation for oral manipulation as they learn instinctual foraging skills (Velasquez-Munoz et al., 2019; Strappini et al., 2021). Through supporting their motivated behaviors (Neave, 2025), environmental enrichment provided to young cattle can influence cognition and affective outcomes. By challenging calves to interact with puzzle boxes, the treatment may have stimulated cognitive processes (like problem-solving and exploration) in addition to enabling the previously mentioned physical behaviors. Although our study did not directly assess cognitive performance, the behavioral patterns we observed are consistent with enriched calves engaging in more varied and active interactions, which may support cognitive development. It is important to note that behaviors within the pen during and outside of experimentation were not assessed but could reveal useful insights into the behavioral development and reaction of calves both exposed to cognitive enrichment and not. Specifically, these assessments could extend to conclusions on broader welfare benefits beyond the immediate puzzle-box use.

#### 5.3.1. Summary

In summary, our data showed that calves readily engaged in a variety of behaviors when provided with puzzle box-type enrichment, and that puzzle boxes supporting cognitive challenge further enhanced behavioral durations compared to controls. By enabling calves to express a wide range of species-specific behaviors, providing the opportunity to access such enrichments can contribute to improved welfare and potentially support cognitive development. Specifically, supporting cognitive functioning through positive experiences can enhance welfare and likely future elements like productivity later in life (Neave, 2025). The fact that both treatments succeeded in facilitating behavioral expression highlights the importance of providing opportunities for calves to engage with their environment and the enhanced behavioral durations with the puzzle boxes emphasizes that more complex interventions may yield additional benefits. These findings encourage further investigation into how cognitive enrichment design can be optimized for calf welfare and behavioral development.

## 6. CONCLUSION

Through our pilot study, we were able to investigate the motivations of calves to voluntarily access and interact with cognitive enrichment when given the choice. We found that calves consistently interacted with all three puzzle box variations and were motivated to visit the enrichment area across all 18 trials regardless of performance. Additionally, we found that calves used a wide range of behavioral expressions to interact directly with the enrichment and the surrounding area, indicative of natural behavior satisfaction. With that in mind, there are still some limitations and unexplored areas of our experiment worth mentioning. Firstly, we did not measure short– and long-term cognitive outcomes, so we cannot confirm that our cognitive enrichment translates into long-term developmental benefits and coping capacities. However, based on previous cognitive research in calves, it is predicted that improved cognitive function early in life can lead to more flexible, adaptable and resilient cows when facing management challenges later in life (Neave, 2025). Thus, even shorter exposures to cognitive enrichment like our own experiment can have long lasting results. Secondly, though we did not consider social factors, it is highly possible that they influenced individual use of cognitive enrichment due to the hierarchal social dynamics of calves housed in groups (Bøe & Færevik, 2003). For instance, it is possible that competition and dominance may have influenced calf participation such that resource guarding may have been present. Thus, we suggest considering social dynamics as a factor moving forward when striving for equal access to cognitive enrichment in a group setting. Thirdly, we believe it would be valuable to apply cognitive enrichment to calves of different age, sex and housing systems to get a broader understanding of the topic under multiple contexts. Lastly, we worked with a small sample size of 10 from the same farm and rearing background, which yielded interesting results but could be limited due to generalizability. Thus, we encourage further studies to build on our findings and contribute new information. There are many different forms of cognitive enrichment, and it would be interesting to see other methods and mechanisms explored to uncover possible interaction variations and individual preferences. With that in mind, further work is still needed to build a stronger understanding of cognitive enrichment for young cattle since this is currently an emerging topic and there are many avenues left to explore.

## FUNDING

This project was supported by Natural Sciences and Engineering Research Council of Canada (NSERC Alliance grant; ID# ALLRP 570894-2021), Prompt, Novalait, Dairy Farmers of Canada, Dairy Farmers of Ontario, Les Producteurs de lait du Québec, and Lactanet, through the Vasseur and Diallo Research and Innovation Chair in Animal Welfare and Artificial Intelligence (WELL-E). Additional stipend funding was generously provided by the Elizabeth and Andre Rossinger Fellowship, and the McGill Graduate Excellence reward.

## CREDIT AUTHORSHIP CONTRIBUTION STATEMENT

**Georgiana Amarioarei**: Conceptualization, Data Curation, Formal Analysis, Investigation, Methodology, Visualization, Writing – Original Draft, Writing – Review & Editing. **Marjorie Cellier**: Conceptualization, Formal Analysis, Investigation, Methodology, Supervision, Visualization, Writing – Review & Editing. **Nadège Aigueperse**: Conceptualization, Supervision, Writing – Review & Editing. **Tania Wolfe:** Conceptualization, Formal Analysis, Investigation, Methodology, Supervision, Writing – Review & Editing. **Elise Shepley**: Conceptualization, Writing – Review & Editing. **Abdoulaye B. Diallo**: Funding Acquisition, Project Administration, Supervision, Writing – Review & Editing. **Elsa Vasseur**: Conceptualization, Funding Acquisition, Methodology, Project Administration, Supervision, Writing – Review & Editing.

## Notes

### Competing Interest Statement

The authors have declared no competing interest.

