## Supplemental Materials for "Investigating cognitive enrichment for dairy calves through behavioral measures of participation and engagement: a pilot study"

**Table S1**. Results from the pairwise comparison of behavior frequencies between puzzle box treatment and control box groups.

| Behavior | Estimate | p-value | t-ratio |
|---|---|---|---|
| Biting | 2.98 ± 2.73 | 0.30 | 1.09 |
| Eating | -3.72 ± 2.73 | 0.20 | -1.36 |
| Environmental interaction | -1.69 ± 2.73 | 0.55 | -0.62 |
| Head/neck rubbing | 0.61 ± 2.73 | 0.82 | 0.22 |
| Licking | 4.02 ± 2.73 | 0.17 | 1.47 |
| Pushing | 3.49 ± 2.73 | 0.23 | 1.28 |
| Sniffing | 2.98 ± 2.73 | 0.30 | 1.09 |
| Standing/walking | -4.10 ± 2.73 | 0.16 | -1.50 |
| Withdrawal | -0.066 ± 2.73 | 0.98 | -0.02 |

**Table S2**. Results from the pairwise comparison of behavioral durations between puzzle box treatment and control box groups.

| Behavior | Estimate | p-value | t-ratio |
|---|---|---|---|
| Biting | 1.26 ± 0.44 | 0.013 | 2.86 |
| Eating | -0.90 ± 0.44 | 0.063 | -2.03 |
| Environmental interaction | -0.12 ± 0.44 | 0.78 | -0.28 |
| Head/neck rubbing | 0.65 ± 0.44 | 0.16 | 1.47 |
| Licking | 0.81 ± 0.44 | 0.090 | 1.83 |
| Pushing | 1.33 ± 0.44 | 0.0099 | 3.01 |
| Sniffing | 0.66 ± 0.44 | 0.16 | 1.49 |
| Standing/walking | -0.07 ± 0.44 | 0.88 | -0.16 |
